# Spatio-temporal control of myoblast identity drives muscle diversity in the Drosophila leg

**DOI:** 10.1101/2025.09.10.675298

**Authors:** Camille Guillermin, Violaine Tribollet, Mathilde Bouchet, Anne Laurencon, Dan Zhou, Sergio Sarnataro, Laurent Gilquin, Isabelle Stevant, Guillaume Marcy, Emeric Texeraud, Yad Ghavi-Helm, Benjamin Gillet, Sandrine Hughes, Samantha Vonau, Jonathan Enriquez

## Abstract

Skeletal muscles display remarkable morphological diversity, but the developmental mechanisms specifying distinct muscle morphology remain poorly understood. Using Drosophila leg muscles as a model, we uncover how naïve mesodermal precursors progressively acquire lineage-restricted identities through a stepwise specification program guided by epithelial morphogens.

Initially, multipotent mesodermal precursors become spatially and transcriptionally restricted into two broad lineages - proximal and distal - under the combined influence of Wg/Wnt1 and Dpp/BMP signals from the overlying epithelium.

By tracing mesodermal precursors that eventually give rise to distal leg muscles, we reveal a second sequence of fate bifurcations that generate distinct muscle subtypes prior to myoblast fusion, as well as a separate lineage producing neuronal lamella cells. Focusing on a single muscle lineage, we show that Wg and Dpp act again during a second phase to control the spatial and temporal deployment of specific transcription factors, ultimately specifying a unique muscle identity.

These findings demonstrate that epithelial morphogens not only pattern the epithelium but also orchestrate muscle diversity by promoting stepwise mesodermal specification. Mesodermal precursors translate morphogen signals over time and space to activate distinct transcriptional programs that operate in parallel with the general program of myogenesis, enabling the emergence of distinct muscle and non-muscle lineages whose unique identities underpin their specialized functions.

## INTRODUCTION

What sets muscles apart from other organs is that, while they all share the property of contractility, they exhibit remarkable morphological diversity. This wide range of muscle morphologies found across the animal kingdom enables complex movements that are essential for functions such as feeding, mating, and even sophisticated tasks like writing articles. One striking example of this diversity are is the muscles located in appendages, which many animals use for various behaviors, including locomotion. For example, human limbs and arms contain around 25 muscles^1^, a number comparable to the distinct muscles found in *Drosophila* legs^2,3^, allowing movement of their appendages in all directions of space. Muscles are also unique in terms of development, as they are composed of syncytia called multinucleated myofibers^4^. These fibers originate from the fusion of specific cell types called myocytes in vertebrates. Despite extensive research into the general mechanisms of muscle development (*i.e.*, how myoblasts generate muscles), the molecular mechanisms controlling the diversity of muscle morphologies in multifiber muscles (*i.e.*, how myoblasts produce distinct muscle types) remain poorly understood.

In vertebrates, muscle progenitor cells (MPCs) originate from the hypaxial region of the dermomyotome, where they delaminate and migrate to populate the limb bud^5^. Once within the limb bud, MPCs undergo active proliferation and become determined to muscle production. These myoblasts then differentiate into myocytes, which fuse to form multinucleated fibers. As this differentiation progresses, myofibers align to form muscle bundles, which will later develop into the muscles^6^. The current model suggests that MPCs are naïve and build individual muscles passively by following a pre-pattern established by muscle connective tissue (MCT)^6–9^.

The progression of myogenesis is tightly regulated by a hierarchical cascade of transcription factors (TFs) that are expressed at different stages of development. During delamination, muscle progenitors express TFs such as Pax3 and Pax7^10,11^, while during migration, they express TFs like Lbx1 and Mox2/3^12,13^. It is only during the proliferative phase that muscle precursors express Myogenic Regulator factors (MRFs)^14^, which are essential for muscle specification and differentiation. Among the MRFs, MyoD and Myf5 are expressed during the proliferation and determination of myoblasts, while Myogenin, Myf6, and MyoD are involved in myocyte differentiation. Additionally, another critical group of TFs in the broader myogenesis program includes members of the Mef2 family, which act downstream and synergistically with the MRFs to coordinate the complex process of muscle development^15^. We hypothesize that myoblasts may generate distinct muscle types, not passively but actively, by expressing specific gene networks in parallel with the general myogenic program.

In *Drosophila*, leg muscle architecture and development closely mirror patterns observed in vertebrate limbs. The adult *Drosophila* leg is structured into segments containing multifiber muscles^6^. Adult muscle precursors (AMPs)^16^ originate from the mesoderm and colonize the leg disc, which is analogous to the limb bud in vertebrates. Within the leg disc, AMPs undergo a phase of proliferation during the larval stages. In early pupal stages, myoblasts fuse to form the adult leg muscles^6^. The transcriptional control of myogenesis is relatively simpler in *Drosophila*. The analog of MRFs in *Drosophila* is the TF Twist (Twi), which can induce myogenesis when overexpressed in embryonic epithelial cells^17^. Although this has not been demonstrated in leg myoblasts, Twi is known to regulate the expression of Mef2 in embryonic and wing myoblasts^18^. Do myoblasts, as suggested in vertebrates, passively build unique muscles, or is there, in parallel with the general myogenic program, a specific program that controls muscle morphology?

Finally, myoblasts and muscles must coordinate their development with other cell types to form a functional leg. In vertebrates, the model of naïve myoblasts passively colonizing a pre-pattern established by the connective tissue offers a simple mechanism of inter-tissue communication during development. However, it remains possible that this pre-patterning - or signals from other surrounding tissues - could actively influence myoblast specification, transitioning them from a naïve state to a specific muscle lineage. In *Drosophila*, external cues from the wing disc epithelium have been shown to pattern indirect and direct flight muscles, leading to the formation of two types of muscle populations with distinct sarcomeric organization and contractile property^19–21^. These studies raise the possibility that epithelial-derived morphogens may not only be instructive for epithelial patterning, but also act as a driving force in myoblast specification, guiding the transition from a naïve state to defined muscle lineages prior to the fusion process.

Here, we combined lineage tracing, single-cell transcriptomics, and spatial imaging to reconstruct the developmental trajectory of mesodermal precursors (MPs) in the Drosophila leg. We identified previously unrecognized muscles and showed that MPs progressively acquire distinct spatial identities through a multistep process that culminates in diverse muscle lineages. Initially naïve, MPs become specified by mid-larval stages into two major lineages: distal muscles in the telopodites and proximal muscles in the prothorax and coxopodite. By following the lineage that generates distal muscles, we found that it undergoes successive fate bifurcations during late larval and early pupal stages, giving rise to three muscle lineages - each producing distinct muscle subpopulations or individual muscles - and one non-muscle lineage that produces neural lamella cells. This diversification is coordinated by positional cues from the leg disc epithelium and involves the progressive restriction of transcriptional profiles within these lineages. Notably, each muscle lineage gives rise to groups of myoblasts characterized by distinct combinations of TFs, expressed in parallel to core myogenic regulators, that are essential for specifying unique muscle identities. Our findings reveal how spatial patterning and lineage dynamics shape the emergence of muscle diversity in the Drosophila leg and provide a framework to study the evolution of appendicular muscle patterning.

## RESULTS

### Diversity of muscle morphologies in the T1 thoracic segment

The muscle architecture of the T1 leg has been documented in previous studies^2,3^. Here, we reexamined the architecture of leg muscles, including those in the thorax responsible for prothoracic leg movement, using a new protocol (see Materials and Methods) we identified three previously undocumented muscles: the coxa levator muscle 2 (*clm2*) in the prothorax, the trochanter reductor muscle 2 (*trrm2*) in the coxa, and the tibial depressor muscle (*tidm2*) in the femur (**Fig. 1A1-A4, Fig. S1 and Fig. S2, Video 1)**.

### Description of mesodermal precursors and myoblasts behavior during development

In our quest to understand how muscle diversity arises during development, we conducted a detailed analysis of Adult Muscle Precursors (AMPs) behavior **(Fig. 1)**. Based on our findings **(Fig. 4)**, we have reclassified the cells previously identified as AMPs as mesodermal precursors (MPs), since these cells contribute to more than just muscle formation.

**Figure 1.**
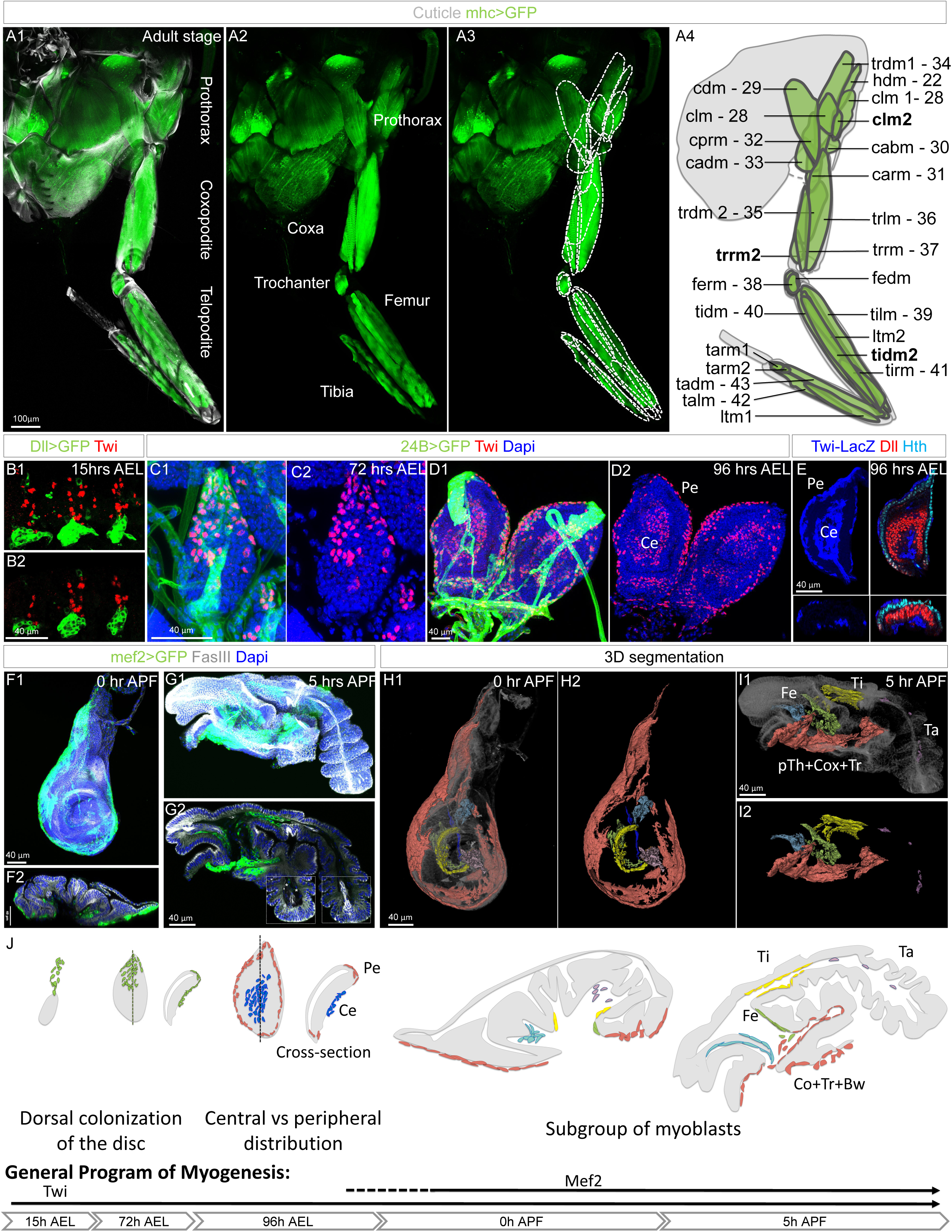
Adult muscle diversity and the progressive spatial organization of muscle progenitors (MPs) into specific clusters. **(A1–A4)** Maximum projection of an adult leg muscles genetically labeled with *UAS-mCD8::GFP* (green) under the control of *MHC-Gal4* **(A1–A3)**, and a schematic representation of all adult muscles **(A4)**. Muscle nomenclature follows Soler and Laurence articles^2,3^: muscle; hd: head dorsal; c: coxa; tr: trochanter; fe: femur; ti: tibia; ta: tarsal; l: levator; d: depressor; ab: abductor; ar: anterior rotator; pr: posterior rotator; ad: adductor; r: reductor; lt: long tendon or Miller’s nomenclature^48^: numbers. **See also Fig. S1 and S2, Video 1**. **(B1–B2)** Maximum projection **(B1)** and confocal section **(B2)** of three thoracic hemi-segments of a stage 16 embryo. Epithelial cells of the leg disc are genetically labeled with *UAS-mCD8::GFP* (green) under the control of *Dll-Gal4*, and MPs are immunolabeled with anti-Twist (red). **(C1–C2)** Maximum projection of a T1 leg disc at 72 hours AEL, with MPs labeled with *UAS-mCD8::GFP* (green) under the control of *24B-Gal4* and stained with anti-Twist (red); nuclei are counterstained with DAPI (blue). **(D1–D2)** Same as **(C1–C2)**, but at 96 hours AEL. **(E)** T1 leg disc at 96 hours AEL carrying *Twi-LacZ* (blue). Shown are a maximum projection (top left), a confocal section (top right), and cross-sections (bottom). Epithelial cells are stained with anti-Dll (red) and anti-Hth (cyan). *Note: Twi-LacZ*⁺ MPs are localized centrally (noted as Ce is the figure) beneath Dll⁺ epithelial cells, and peripherally (noted as Pe is the figure) adjacent to Hth⁺ epithelial cells. **(F1–G2)** T1 discs at pupal stages: 0 hours APF **(F1–F2)** and 5 hours APF **(G1–G2)**. MPs are genetically labeled with *UAS-mCD8::GFP* (green) under the control of *Mef2-Gal4*, stained with anti-Fas3 (gray), and nuclei counterstained with DAPI (blue). F1 and G1 show maximum projections; F2 and G2 show cross-sections. The dotted boxes highlight the weak GFP expression observed in the tarsus (asterisks), visible in a different focal plane. **(H1–I2)** 3D segmentation of MPs from the discs shown in (F1–G2). pTh: Prothorax; Cox: Coxa; Tr: trochanter; Fe: Femur; Ti: Tibia; Ta: Tarsus. **See also Video 2**. **(J) Top:** Schematic of the coordinate development of the leg MPs/myoblasts and the epithelium. Below: the schematic of the general program of myogenesis. **See also Fig.S3.**

Our study begins at the late embryonic stage, when the epithelial cells of the leg disc have evaginated from the ectoderm (stage 16, 14 hours After Egg Laying [AEL]). At this stage, approximately 20 mesodermal precursors (Twit^+^) are closely associated with the dorsal region of epithelial precursors of the leg discs (Dll^−^ GFP^+^) **(Fig. 1B, Fig. S3)**. As development progresses, by 72 hours AEL, the number of MPs increases to around 50. At this stage, MPs also begin to colonize the dorsal region of the disc **(Fig. 1C, Fig. S3)**. By 96 hours AEL, the number of MPs increases dramatically to approximately 450, fully colonizing the leg imaginal disc and forming two distinct MP clusters: central and peripheral **(Fig. 1D, Fig. S3)**.

During late larval stages, Mef2 expression initiates in a subset of Twi⁺ cells, particularly at the periphery **(Fig. S3)**. By 0 hours After Pupal Formation (APF), all Twi⁺ cells express Mef2>GFP **(Fig. S3).** However, we observed a small population of cells with low Mef2>GFP levels and very low or undetectable Twi expression, particularly in the tarsus, suggesting the specification of some MPs toward a non-muscle fate **(see subsequent sections, Fig. 3 and Fig. S4)**. At this stage, the central myoblast cluster subdivides into four smaller clusters, each localizing to specific presumptive territories: two in the femur, one in the tibia, and one in the tarsus **(Fig. 1F, H; Video 2)**. By 5 hours APF, evagination occurs in the tarsal and tibial segments, with partial protrusion of the femoral segment. Notably, the cluster in the tarsal segment exhibits very low Mef2>GFP signal **(Fig. 1G2)** and no detectable Twi protein **(Fig. S3)**, supporting the specification of these myoblasts toward a non-muscle lineage **(see subsequent sections, Fig.3 and Fig. S4)**. Meanwhile, the coxa, trochanter, and prothoracic segments remain un-evaginated, with the myoblasts in these regions organized into two concentric rings **(Fig. 1H1-I2).**

**Figure 2.**
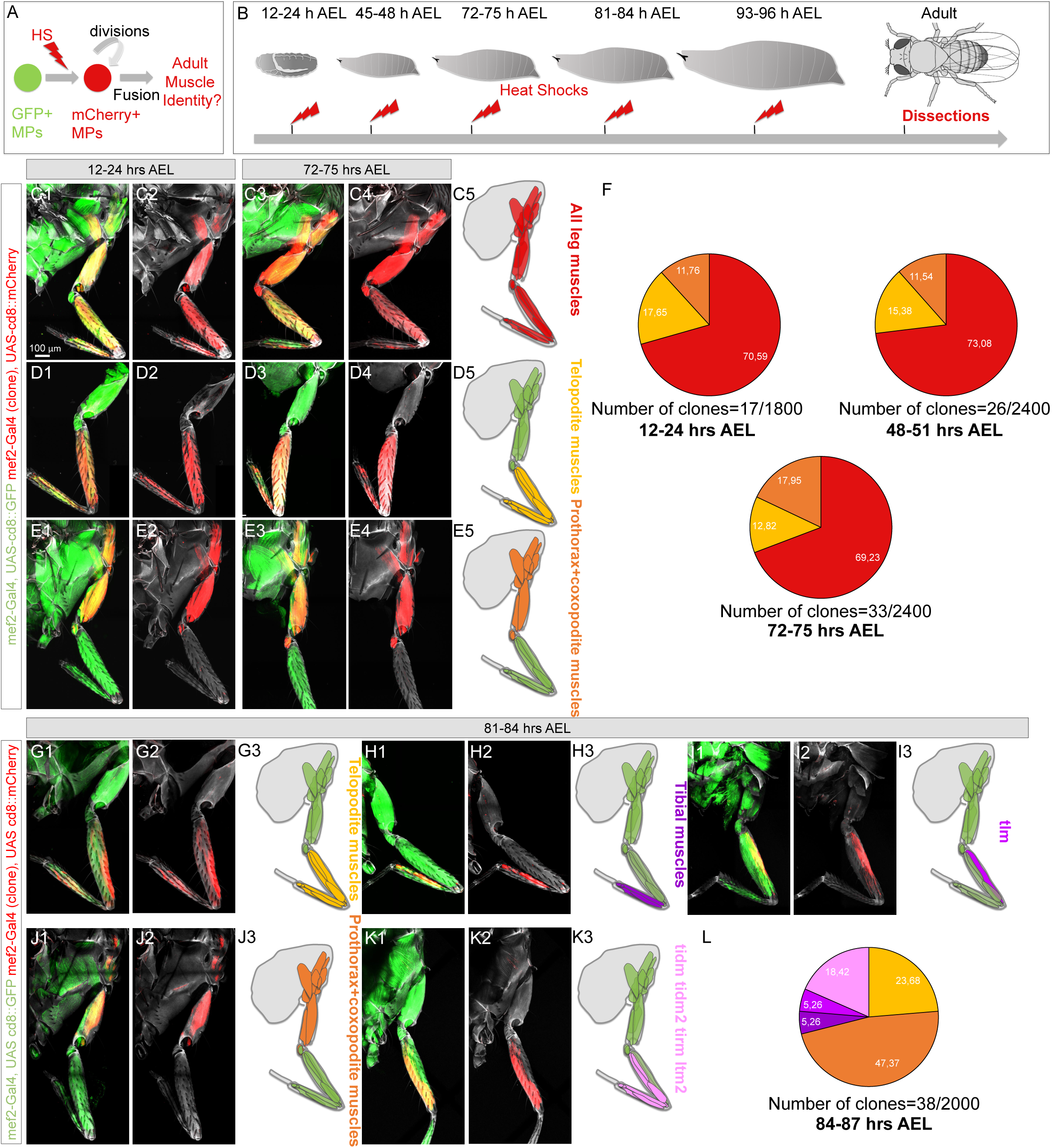
Random lineage tracing of MPs. **(A)** Schematic of the clonal induction of mCherry in MPs. **(B)** Schematic of the timing of the clonal induction of mCherry in MPs. **(C1-E5)** Maximum projection and schematic of the T1 leg and prothorax where all muscle are labeled with *UAS-mCD8::GFP* (green) under the control of *Mef2-Gal4* showing the expression of mCherry in muscle after its induction in MPs at 12-24 hours or at 72-75 hours AEL. The expression of the mCherry expression is color coded in the schematics of the leg. Note: we used the FB1.1 system to induce randomly MP clones, (**See Material & Methods). See Fig. S4** **(F)** Circular diagram showing the portion of the three types of clones found after mCherry induction at 84-87 hours AEL. **See Fig. S4** **(G-L)** Confocal images and corresponding schematics of the T1 leg where clones’ induction in MPs is triggered at 81-84 hours AEL. mCherry expression is color coded in the schematics, and the respective proportions are presented on a pie chart.

**Figure 3.**
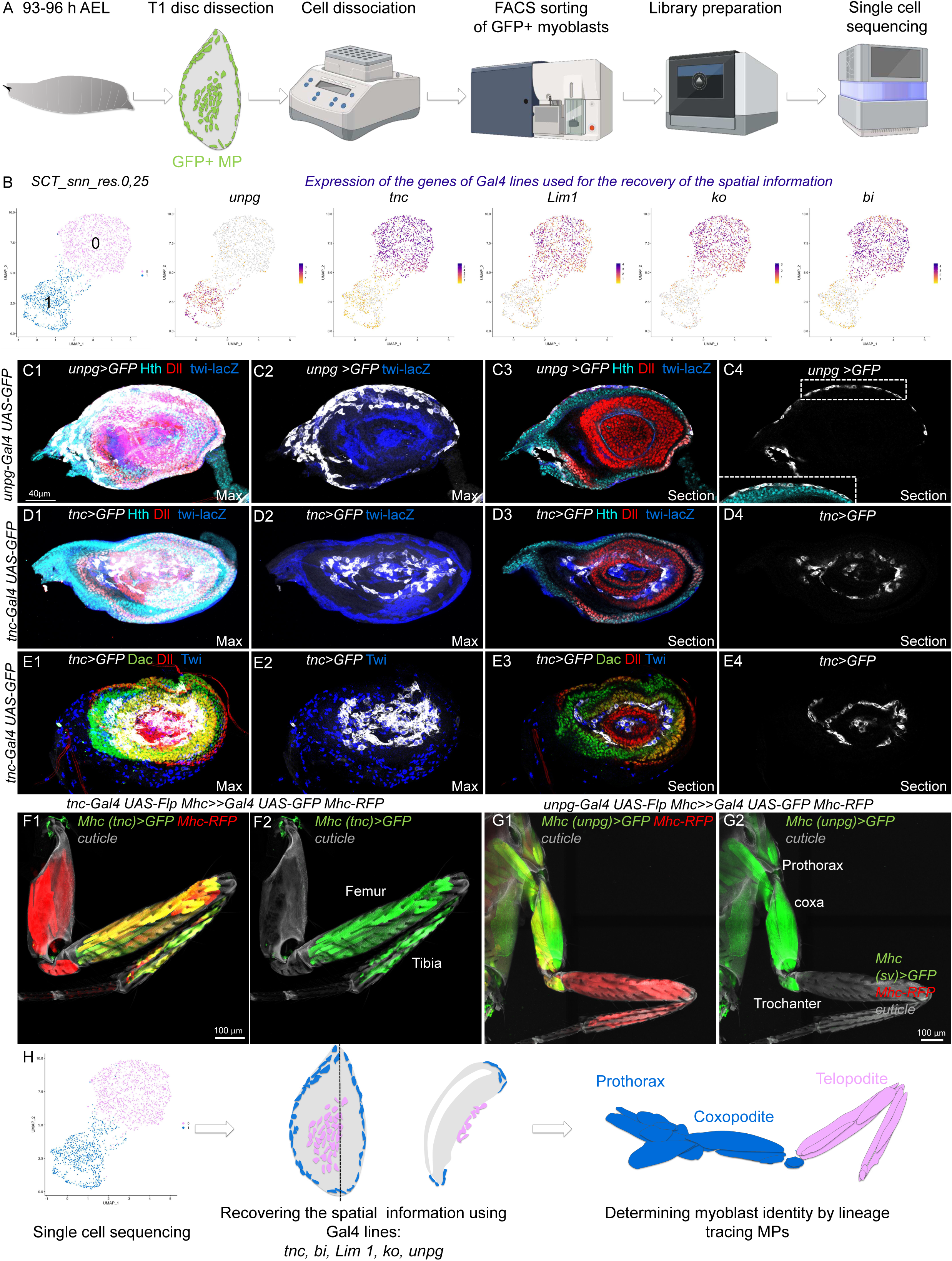
Two spatially and transcriptionally distinct populations of MPs give rise to proximal and distal muscles. **(A)** Schematic of the scRNA-seq workflow. **(B)** UMAPs showing two MP clusters (left) and the expression of spatial identity genes used in panels (**C1–E4)** (right). See also Fig. S5. **(C1–E4)** Leg discs (96 hours AEL) from *Twi-lacZ* (blue) larvae genetically labeled with *UAS-mCD8::GFP* under the control of *unpg-Gal4* **(C1–C4)** or *tnc-Gal4* **(D1–E4)**, and immunostained with anti-Dll (red) and anti-Hth (cyan) **(C1–D4)**, or with anti-Dll (red) and anti-Dac (green) **(E1–E4)**. The dotted boxes in (C4) highlight the expression of Hth in peripheral unpg-GFP+ MPs. **(F1–G2)** T1 legs labeled with *Mhc-RFP* (red), showing GFP expression in distal and proximal muscles when the lineage tracing is activated in MPs using *tnc-Gal4* **(F1–F2)** or *unpg-Gal4* **(G1–G2)**, respectively. **(H)** Schematic summarizing the workflow used to determine the spatial position and identity of MP clusters.

**Figure 4.**
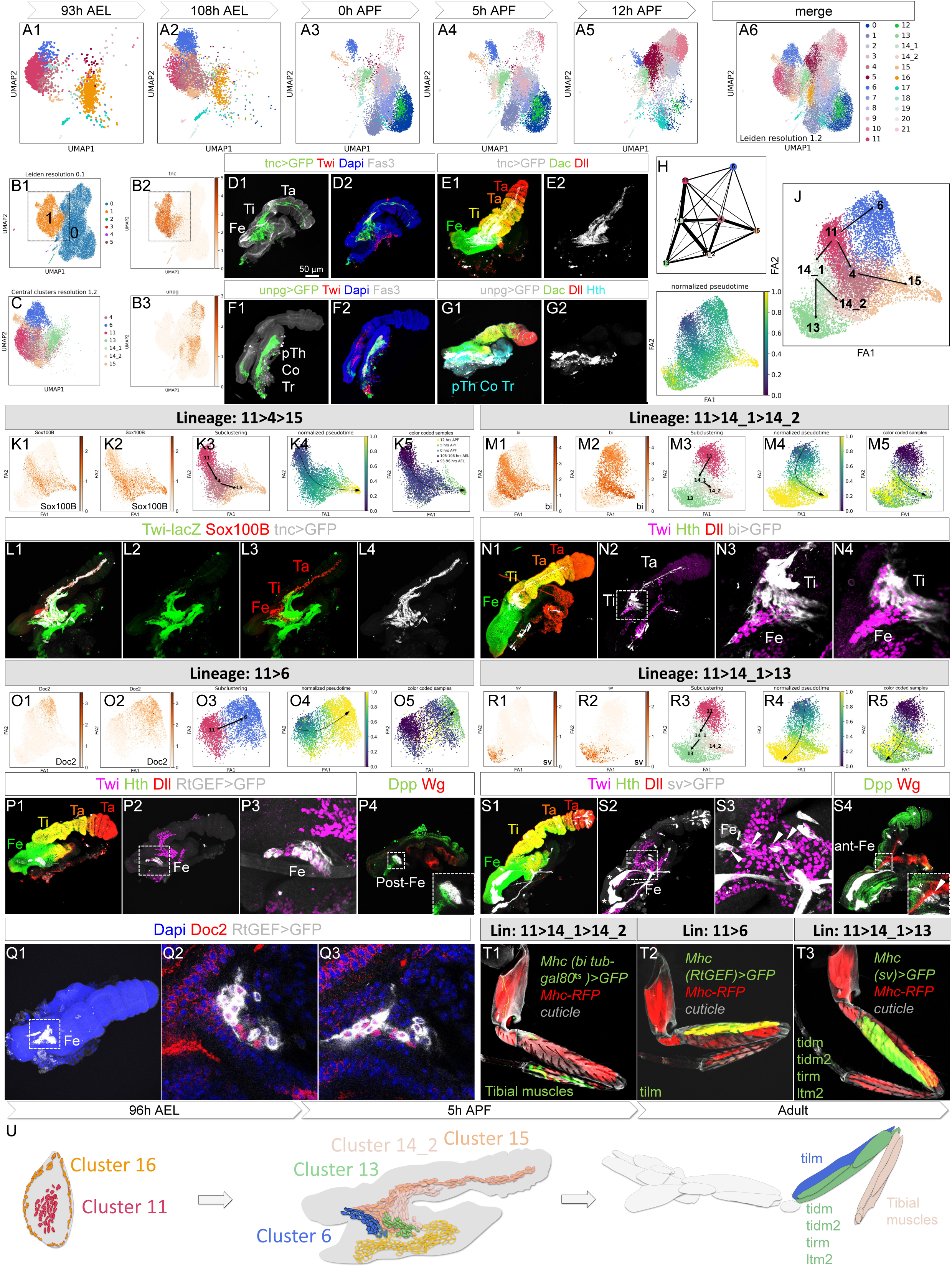
The central cluster of MPs gives rise to four spatially and transcriptionally distinct lineages. **(A1–A5)** UMAPs of GFP^+^ MPs FACS-sorted at different time points **(A1–A5)** and a merged UMAP of all time points. **(B1-B3)** Low-resolution UMAP revealing two major clusters **(B1)** and the Differential expression of *tnc* **(B1)** and *unpg* **(B1) in these two clusters**. **(C)** high-resolution UMAP of cluster 1 (central cluster) in **(B1)**. **(D1–G2)** T1 imaginal discs at 5 hours APF genetically labeled with *UAS-mCD8::GFP* (green or white) under the control of *tnc-Gal4* **(D1–E2)** or *unpg-Gal4* **(F1–G2),** immunostained with Fas3 (grey), Twi (red), and DAPI (nuclei, blue) **(D1–D2, F1–F2),** or with Dac (green), Dll (red), and Hth (cyan) **(E1–E2, G1–G2)**. **(H–I)** PAGA connectivity tree **(H)** and ForceAtlas2 (FA2) map color-coded by pseudotime **(I)** of the central cluster in **(C)**. **(J)** FA2 map showing distinct clusters. Arrows indicate the selected lineages based on the PAGA tree **(H)**, FA2 pseudotime **(I, K4, M4, O4, R4)**, and real-time FA2 projection. **(K1–R5)** FA2 maps showing the expression of different lineage marker of the central clusters **(K1, M1, O1 and R1)** and of individual lineages **(K2, M2, O2, R2); (K3-K5, M2-M5, O2-O5, R2-R5)** are FA2 maps of individual lineages where cluster are color coded, pseudotime or real color code time are represented. The expression of markers used to determine lineage identity in immature T1 legs at 5 hours APF: **(L1–L4)** Sox100B (red), *Twi-lacZ* (green), *tnc-Gal4 > UAS-mCD8::GFP* (grey); **(N1–N4)** *bi-Gal4 > UAS-mCD8::GFP* (grey), Dll (red), Hth (green), Twi (purple); **(P1–Q3)** *RtGEF-Gal4 > UAS-mCD8::GFP* (grey) with Dll (red), Hth (green), Twi (purple) **(P1–P3)**, Dpp (green) and Wg (red) **(P4)**, or Doc2 (red) and DAPI (blue) **(Q3)**; **(S1–S4)** *sv-Gal4 > UAS-mCD8::GFP* (grey) with Dll (red), Dac (green), Twi (purple) **(S1–S3),** or Dpp (green) and Wg (red) **(S4)**. **(T1–T3)** T1 legs labeled with *MHC-RFP* (red), showing GFP expression in distal and proximal muscles following lineage tracing from *bi-Gal4* **(T1)**, *RtGEF-Gal4* **(T2)**, or *sv-Gal4* **(T3)**. Note: *tubP-GAL80<u>^ts^</u>* was used to activate lineage tracing only after 0 hours APF. Asterisks indicate weak GFP expression in a muscle located in the tibia **(U)** Schematic summarizing the spatial localization of scRNA-seq clusters in 5 hours APF immature legs and the corresponding adult muscles.

In summary, MPs colonize the disc dorsally, initially organizing into central and peripheral groups. The central cluster subsequently subdivides into four smaller clusters, suggesting that this spatial organization may prefigure the future arrangement of muscles or other cell types **(Fig. 1J)**. This spatial pattern of behavior prior to fusion closely resembles that observed in vertebrate limb muscle precursors.

### MPs are progressively committed to producing specific muscles

To assess whether MPs are specified to produce distinct muscles before fusion, we generated clones at different developmental stages **(Fig. 2A-B)**. Previous studies^3^ suggested that, during early larval stages, MPs have the potential to generate all prothoracic muscles (the legs were not analyzed, as they had been removed in the experiment).

To determine whether MPs are specified before fusion, we first generated mCherry^+^ clones during the second half of embryogenesis, 12-24 hours AEL. Among 900 examined flies, we identified 17 clones in the prothoracic segment. The majority of these clones (**70%, N=12/17**) labeled all leg muscles across all leg segments, including the prothorax, a proportion consistent with our observations in the mesothoracic and metathoracic segments, though not detailed here (**Fig. 2C1-C2, C5, F**).

**Figure 5.**
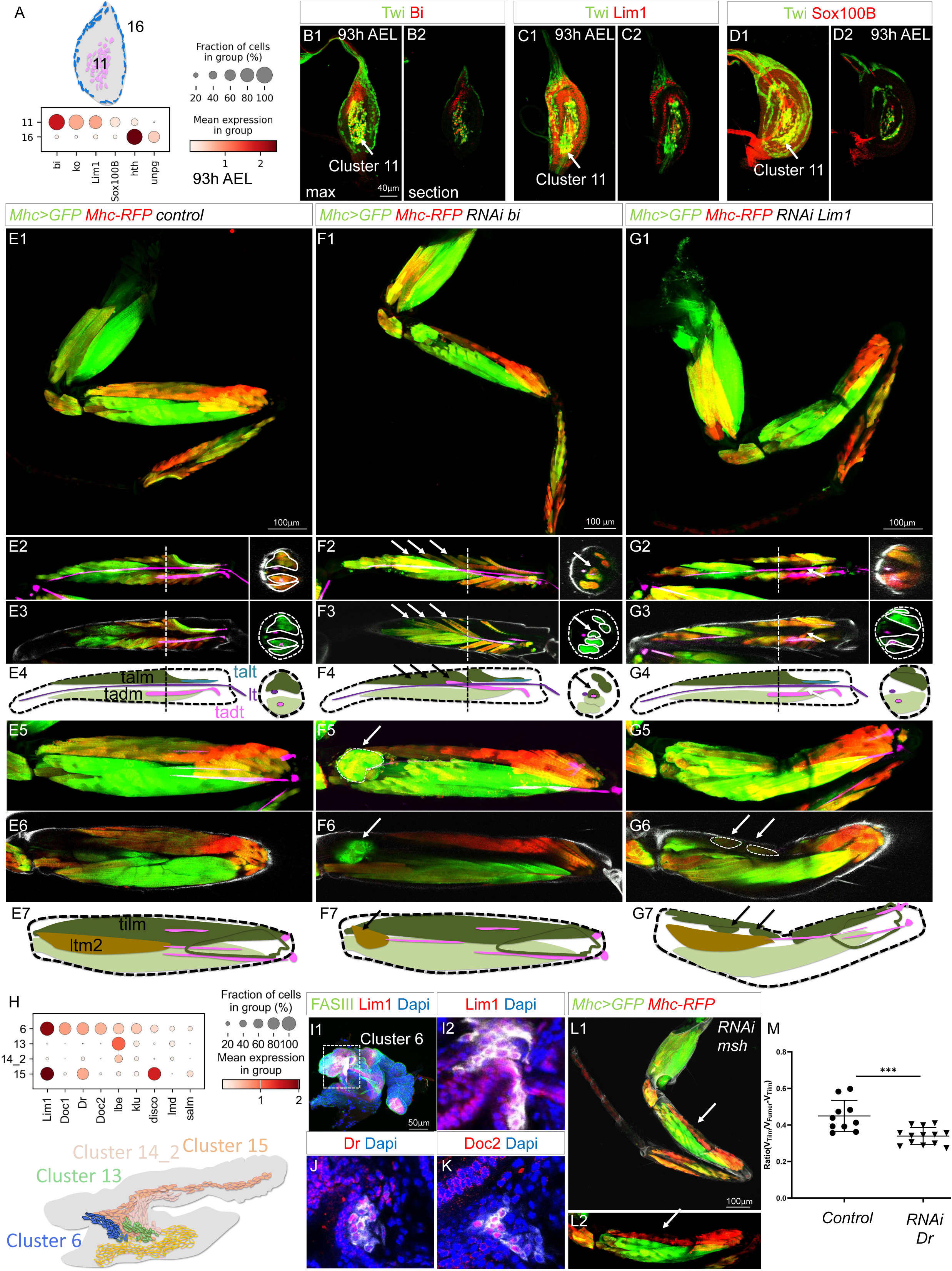
Muscle diversity is controlled by distinct combinations of transcription factors. **(A)** Dot plot showing the expression of RNAs coding for TFs differentially expressed in MPs of cluster 11 *vs.* cluster 16 at 93-96 h AEL. **(B1–D2)** T1 leg discs at 93-96 hours AEL immunostained for Twi (green) and Bi (red) (**B1–B2)**, Lim1 (red) **(C1–C2)**, or Sox100B (red) **(D1–D2)**. Panels **(B1, C1, D1)** show maximum projections; **(B2, C2, D2)** show single confocal sections. **(E1–G7)** T1 legs genetically labeled with *MHC-RFP* (red) and *UAS-mCD8::GFP* (green) under the control of MHC-Gal4 in a control fly **(E1–E4)**, a fly with *bi* knockdown (KD) in MPs, myoblasts, and muscles **(F1–F7)**, a fly with *Lim1* KD in MPs, myoblasts, and muscles **(G1–G7)**. Note: The two MHC enhancers driving RFP and Gal4 expression have different sequences and display differential muscle activity, allowing us to better distinguish them. **(E2–E4, F2–F4, G2–G4)** Enlargements of the tibia: **(E2, F2, G2)** are maximum projections; **(E3, F3, G3)** are confocal sections; **(E4, F4, G4)** are schematics of the tibia showing the talm, tadm, and their corresponding tendons, talt (cyan) and tadt (pink); the long tendon (lt) is also represented (purple). Cross sections on the right are indicated by dotted lines on the left. Note: In **(F2–F4)**, the talm attaches to the wrong tendon (tadt, arrows in **F2-F4**) when bi is KD in MPs, myoblasts, and muscles. In **(G2-G4)**, arrows highlight fibers occasionally misattached to tendons when Lim1 is KD in MPs. **(E5–E7, F5–F7, G5–G7)** Enlargements of the femur: **(E5, F5, G5)** are maximum projections; **(E6, F6, G6)** are confocal sections; (**E7, F7, G7)** are schematics of the femur showing four femoral muscles and their tendons (pink). Note: In **(F5–F7)**, ltm2 appears rounded and retracted (arrow) while still attached to its tendons when *bi* is KD in MPs, myoblasts, and muscles. In **(G5–G7)**, the leg is folded and proximal tilm fibers are rounded and appear detached from tendons (arrow in **G6-G7**) when *Lim1* is KD in in MPs, myoblasts, and muscles. Tendons are labeled in purple (see Material & Methods). **(H)** Dot plot showing the expression of RNAs coding for TFs differentially expressed in myoblasts of cluster 6 vs. central clusters (13, 14_2, 15) at 5 hours APF. **(I1–K)** Immature legs at 5 hours APF genetically labeled with *RtGEF-Gal4 > UAS-mCD8::GFP* (gray) and immunostained for Lim1 (red, I1–I2), Dr (red, J), or Doc2 (red, K). **(I1)** is a maximum projection; **(I2-K)** are confocal sections showing *RtGEF*>GFP+ myoblasts in the presumptive femur territory. **(L1–L2)** T1 legs labeled with MHC-RFP (red) and UAS-mCD8::GFP (green) under MHC-Gal4 in a fly with Dr KD in MPs, myoblasts, and muscles, resulting in an atrophic tilm. **(M)** Graph showing the relative volume of the tilm compared to all femoral muscles in control flies vs. Dr KD flies (in MPs, myoblasts, and muscles).

Interestingly, in a subset of cases (N = 4), clones extended into the contralateral hemisegment, including the leg, an occurrence previously reported in the thorax^3^. Notably, in these instances, muscle labeling intensity was consistently weaker on one side. Given the rarity of such clone events and their absence in the mesothoracic and metathoracic segments, it is unlikely that these observations result from two independent clone events. Instead, we hypothesize that MPs migrate to the contralateral leg disc, a phenomenon that may occur specifically in the prothoracic segment due to physical contact between the T1 leg discs. These findings, where clones label all muscles in the prothoracic segment and can cross the body midline, strongly suggest that mesodermal precursors are naïve prior to colonizing the leg disc.

Additionally, we identified two other distinct MP clone types: one labeling muscles in the femur and tibia (telopodite) (18%, N=3) (**Fig. 2D1-D2, D5, F**), and another labeling muscles in the trochanter, coxa (coxopodite) and prothorax (12%, N=2) (**Fig. 2E1-E2, E5, F**). These findings raise two possible hypotheses: (1) MPs begin to be specified into these two muscle subpopulations during embryogenesis, suggesting that most MPs remain uncommitted; or (2) MPs specification occurs at later developmental stages.

To distinguish between these scenarios, we generated MPs clones during larval stages. Clones induced at 48 hours AEL **(Fig. 2F**) and 72 hours AEL **(Fig. 2C3-C4, D3-D4, E4-E5**) exhibited similar results to embryonically induced clones. However, when clones were induced at 84 hours and 96 hours AEL, we no longer observed clones labeling all muscles **(Fig. 2L)**. Instead, clones predominantly labeled either the tibia and femur **(Fig. 2G1-G3, L)** or the trochanter, coxa, and prothorax **(Fig. 2J1-J3, L)**. Intriguingly, at these later time points, we also identified clones labeling only a subset of muscles within the femur or tibia: (1) clones labeling random subsets of tibial muscles **(Fig. 2H1-H3, L)** (2) clones labeling the tibial levator muscle (*tilm*) in the femur **(Fig. 2I1-I3, L)** and (3) clones labeling the *tidm*, *tirm*, and *ltm2* in the femur **(Fig. 2K1-K3, L)**.

Collectively, these results indicate that MPs enter the disc in an unspecified state and subsequently specify into two populations: one producing distal muscles (tibia and femur) and the other producing proximal muscles (prothorax, coxa, and trochanter) after 82 hours AEL. These two populations may correspond to mesodermal precursors localized in the disc center versus those at the periphery. Furthermore, the presence of subpopulation-specific clones at 84 hours and 96 hours AEL suggests that MPs gradually specify to generate both muscle subpopulations and individual muscles over time.

### The initial step in muscle specification involves defining distal muscles versus proximal muscle lineages

We began by investigating the first step of specification, proximal *vs* distal, by conducting scRNA-seq of GFP^+^ MPs at 96 hours AEL **(Fig. 3A)** and unveiled two distinct cell clusters (clusters 0 and 1) **(Fig. 3B)**.

To recapture the spatial information lost during FACS sorting, we employed GFP expression under the control of enhancer or gene trap lines specifically active in one or the other cluster. In parallel, we labeled the peripheral part of the epithelium, presumptive territory of the prothorax and the coxopodite (coxa and trochanter), and the central part of the epithelium, presumptive territory of the telopodite (femur and tibia) with Hth and Dll/Dac, respectively, to characterize the position of the two MP clusters relative to the epithelium **(Fig. 3C1-E4)**. We observed that MPs of cluster 0 labeled with GFP (*tnc, bi, Lim1, or ko-Gal4* lines) are located at the disc’s center (Dll^+^ and Dac^+^), while GFP^+^ MPs of cluster 1 (*unpg-Gal4* lines) are localized at the periphery of the disc (Hth^+^) **(Fig. 3C1-E4, Fig. S4)**.

We then employed a lineage muscle tracing system to determine the destiny of both MP population until the adult stage. When we activated the lineage muscle tracing system using Gal4 under the control of *tnc*, *bi*, or *ko* (cluster 0), GFP expression was observed in tibial and femoral muscles **(Fig. 3F1-F2, Fig. S4)**. Conversely, activation of the lineage muscle tracing system with *unpg-Gal4* (cluster 1) resulted in GFP expression in muscles of the prothorax, coxa, and trochanter **(Fig. 3G1-G2, Fig. S4)**.

Our clonal analysis, combined with scRNA-seq, revealed that during larval stages, MPs lose their potential to generate all muscle types and instead become specified into two distinct populations: one localized at the center and the other at the periphery of the disc. These populations give rise to distal and proximal muscles, respectively **(Fig. 3H)**. The spatial arrangement of these populations prefigures the future positioning of distal and proximal muscles more than 48 hours before the fusion process, a time during which they exhibit distinct TFs expression profiles **(Fig. 5)**.

### Mesodermal precursors are progressively specified to produce unique muscles

To monitor the progressive specification of MPs, we performed scRNA-seq at various time points on FACS-sorted GFP^+^ MPs (Twi^+^, Mef2^−^) and myoblasts (Twi^+^, Mef2^+^; **Fig. 4A1-A6**). After correcting for batch effects, we pooled the data from each time point and generated a low-resolution UMAP (**Fig. 4B**). This analysis revealed two major clusters that encompass nearly the entire population of MPs/myoblasts.

**Figure 6.**
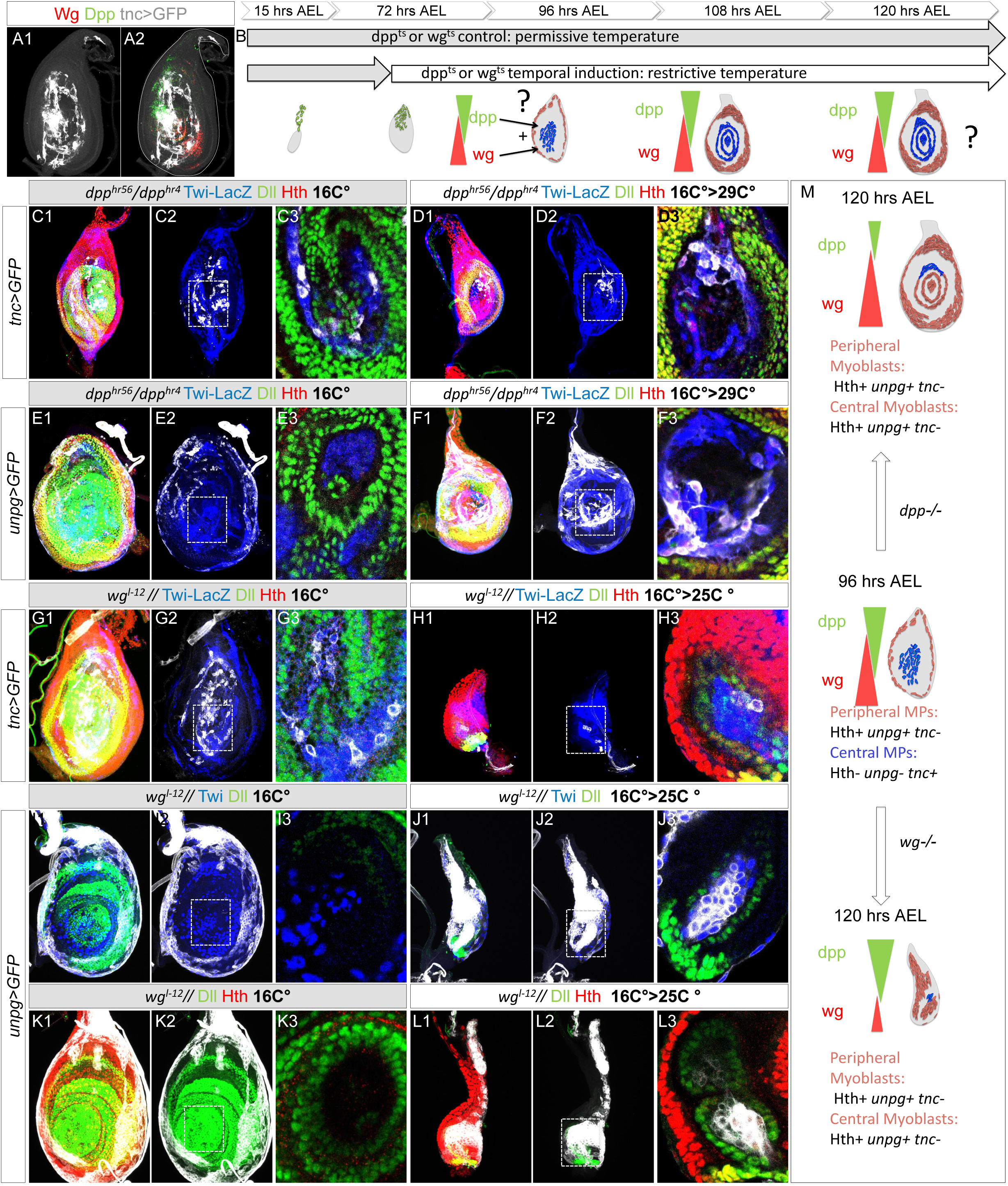
Wg and Dpp act synergistically to specify the central vs peripheral identity of MPs. **(A1-A2)** Late third instar T1 leg disc genetically labeled with *tnc-Gal4>UAS-mCD8::GFP* (gray), immunostained for Dpp (green) and Wg (red). **(B)** Schematic of the conditional *dpp* and *wg* loss-of-function strategy. **(C1-F3)** Third instar leg discs of *Twi-lacZ*; *dpp^hr56^/dpp^hr4^* mutants expressing *tnc-Gal4>UAS-mCD8::GFP* (gray) **(C1–D3)**, or *unpg-Gal4>UAS-mCD8::GFP* (gray) **(E1–F3**), immunostained for Dll (green) and Hth (red). Discs were kept at the permissive temperature **(C1–C3, E1–E3)** or shifted to the non-permissive temperature shortly before the specification of central vs peripheral MPs (**D1-D3, F1-F3)**. **(G1–L3)** Third instar leg discs of *Twi-lacZ*; *wg^1-12^* mutants expressing *tnc-Gal4>UAS-mCD8::GFP* (gray) **(G1**-H3) or *unpg-Gal4>UAS-mCD8::GFP* (gray) **(I1–L3)**. Discs were immunostained for Dll (green) and Hth (red) **(G1-H3, K1-L3)** or for Twi (blue) and Dll (green) **(I1-J3)**. Samples were maintained at the permissive temperature **(G1-G3, I1-I3, K1-K3)** or shifted to the non-permissive temperature before the specification of central vs peripheral MPs **(H1-H3, J1-J3, L1-L3)**. **(M)** Schematic summary of the phenotypic outcomes observed in panels **(C1-L3).**

To recapture spatial information lost during FACS sorting, we used Gal4 drivers (*tnc-Gal4* and *unpg-Gal4*) that are differentially expressed in two major clusters **(Fig. 4C1–C2**). By combining these drivers with morphological (Fas III) and spatial (Dll, Dac, Hth) markers, we determined that *unpg-Gal4* -which predominantly labels MPs/myoblasts in cluster 0-continues to drive GFP expression at later stages in cells located within the presumptive coxopodite and prothoracic territories **(Fig. 4F1-G2)**. In contrast, *tnc-Gal4* -which labels all MPs/myoblasts in cluster 1-drives GFP expression in cells localized to the telopodite territory **(Fig. 4D1-E2)**. These observations indicate that even at late developmental stages, MPs/myoblasts remain spatially and transcriptionally subdivided into central and peripheral populations, suggesting the existence of two distinct specification pathways.

We then focused our investigation on the specification of telopodite-associated MPs/myoblasts (cluster 1). To ensure that these populations correspond to MPs fated to form telopodite muscles even at later stages, we activated the muscle lineage tracing during early pupal stage using *tnc-Gal4* in combination with *tub-Gal80^ts^*. This late induction was still sufficient to label all telopodite muscles, confirming *tnc-Gal4* is a good marker for telopodite lineages even at later stages **(Fig S5)**.

We next generated a higher-resolution UMAP, which revealed six MP/myoblast clusters (4, 6, 11, 13, 14, and 15), and performed a PAGA^22^ trajectory on these central clusters and found four different lineages **(Fig. 4H-J).** This analysis revealed that the central cluster (cluster 11) at 96 hours APF progressively give rise to four muscle lineages **(Fig. 4H-J) (See Material & Methods).** Of note, cluster 14 was subdivided into two subclusters, 14_1 and 14_2, to improve resolution of the lineage trajectories leading to tibial *vs* femoral muscles. To recover spatial information lost during FACS sorting and to determine the identity of these lineages, we combined immunostaining, lineage-specific Gal4 drivers, and our muscle lineage tracing system.

Lineage 11 → 4 → 15 continuously expresses Sox100B **(Fig. 4K1-K5)**. Co-staining with Sox100B and *tnc*-Gal4-driven GFP during early pupal stages revealed tnc⁺/Sox100B⁺ cells in the tarsus, femur, and tibia **(Fig. 4L1-L4)**. Their localization in the tarsus, absence of Mef2, and loss of twist expression at late stages suggest that they are not myoblasts **(Fig. 4L1-L4, Fig. S5)**. Based on their spatial distribution and marker expression confirmed by immunostaining (Lim1⁺, Sox100B⁺) **(Fig. 4L1-L4, Fig. S5)**, as well as by comparing marker profiles in our scRNA-seq dataset with previously published data **(Fig. S5)**, we propose that these cells correspond to the neural lamella cells recently identified in the adult tarsus by scRNA-seq^23^. Our findings extend the presence of this population to the femur and tibia and further demonstrate their mesodermal origin.

The *bi* gene is highly expressed throughout the lineage 11 → 14_1 → 14_2 **(Fig. 4M1-M5)**. A *bi-Gal4* enhancer trap line labels Twi*⁺* MPs localized in the central region of the leg disc at early stages, corresponding to cluster 11, which gives rise to all telopodite muscles **(Fig. S4)**. However, at 0 hours and 5 hours APF, *bi-Gal4*-driven GFP⁺/Twi*⁺* myoblasts (cluster 14_2) are restricted to the presumptive tibia region **(Fig. 4N1-N4; Fig. S5)**. In addition, *bi-Gal4* also labels GFP⁺/Twi*⁻* cells located in the presumptive femur, tibia, and tarsus, corresponding to neural lamella cells (cluster 15), which also express *bi* but at lower level **(Fig. 4M1, Fig. S5)**. To trace the lineage of bi⁺/Twi*⁺* MPs after pupal stages (cluster 14_2), we activated the muscle lineage tracing system at 0 hours APF using *bi-Gal4* in combination with *tub-Gal80^ts^*. This temporally controlled induction specifically labeled tibial muscles, indicating that MPs in cluster 14_2 are committed to the tibial muscle lineage **(Fig. 4T1)**.

To identify lineage 11 → 6, we used *RtGEF-Gal4* to drive GFP expression. We found that this driver initiates expression at the late larval stages (120 hours AEL) and serves as a reliable marker for cluster 6 **(Fig. 4P1-P4, Fig. S5)**, as demonstrated by the expression of the transcription factor Doc2, a specific marker of cluster 6, in RtGEF>GFP^+^ myoblasts **(Fig. 4 O1-P3)**. These Twi^+^ myoblasts are localized in the presumptive territory of the dorsal femur, where the tilm will form **(Fig. 4P1, P4**, **Fig. S5)** and lineage tracing of *RtGEF-Gal4*^+^ myoblasts confirmed that they give rise to the tilm **(Fig. 4T2)**.

For lineage 11 → 14_1 → 13, we used a *sv-Gal4* line expressed in cluster 13 **(Fig. 4R1, R2)**. GFP^+^ myoblasts under the control of *sv-Gal4* were localized in the presumptive territory of the ventral femur only at 5 hours APF which align with their late expression in this lineage as revealed in the in the PAGA graph **(Fig. 4R1, R2, S1-S4, Fig. S5)**. Their lineage tracing showed they give rise to the *ltm2*, *tidm*, *tidm2* and *tirm* **(Fig. 4T3)**.

These results reveal that the central populations of MPs/myoblasts, which give rise to distal leg muscles, are progressively specified into subpopulations characterized by distinct combinations of TFs **(Fig. 5)**. This molecular heterogeneity among MPs/myoblasts underlies the formation of muscle groups and, in some cases, even individual muscles **(Fig. 5)**. Notably, we show that muscle identity is specified early in development -more than 24 hours before the onset of fusion-for myoblasts in cluster 6, which give rise to the *tilm*. This cluster is already spatially restricted to the future location of the adult muscle and appears uniquely capable of fusing only with cells of the same cluster, suggesting an early, tightly regulated lineage-restricted identity.

### Unique TF codes control muscle diversification

At 96 hours AEL, central and peripheral MPs express distinct combinations of TFs **(Fig. 5A)**, which may play key roles in their respective fate specification. We first confirmed the presence of several TFs at the protein level using available antibodies, validating the expression of peripheral TFs such as Hth **(Fig. 3C4)**, and central TFs including Lim1, Sox100B, and Bi **(Fig. 5B1-D2)**.

To assess the functional relevance of the central TFs code, we knocked down two TFs -*Lim1* and *bi*-specifically expressed in central MPs, using *Zfh1-Gal4*, a strong driver active in MPs and myoblasts in the leg disc **(Fig S6)**. Strikingly, only distal muscles (in the femur and tibia) were affected, revealing an instructive role for these TFs in specifying distal muscle identities **(Fig. 5)**.

When *bi* is knocked down in MPs and muscles, some femoral muscles are mildly affected -particularly *ltm2*-but the strongest phenotypes are observed in tibial muscles, which normally maintain *bi* expression during pupal stages **(Fig. 5 F1, F5-F7, Table 1)**. For instance, *tarm1* is occasionally absent or reduced in fiber number **(Fig S6)**, while the *talm* muscle is consistently affected. In some cases, *talm* fibers appear rounded and fail to attach to their correct distal tendon in adults **(Fig S6)**. However, the most consistent phenotype involves *talm* fibers misattaching to the *tadm* tendon (tadt), pulling this tendon toward the center of the tibia near the long tendon *(lt)* **(Fig. 5F1-F4).**

**Figure 7.**
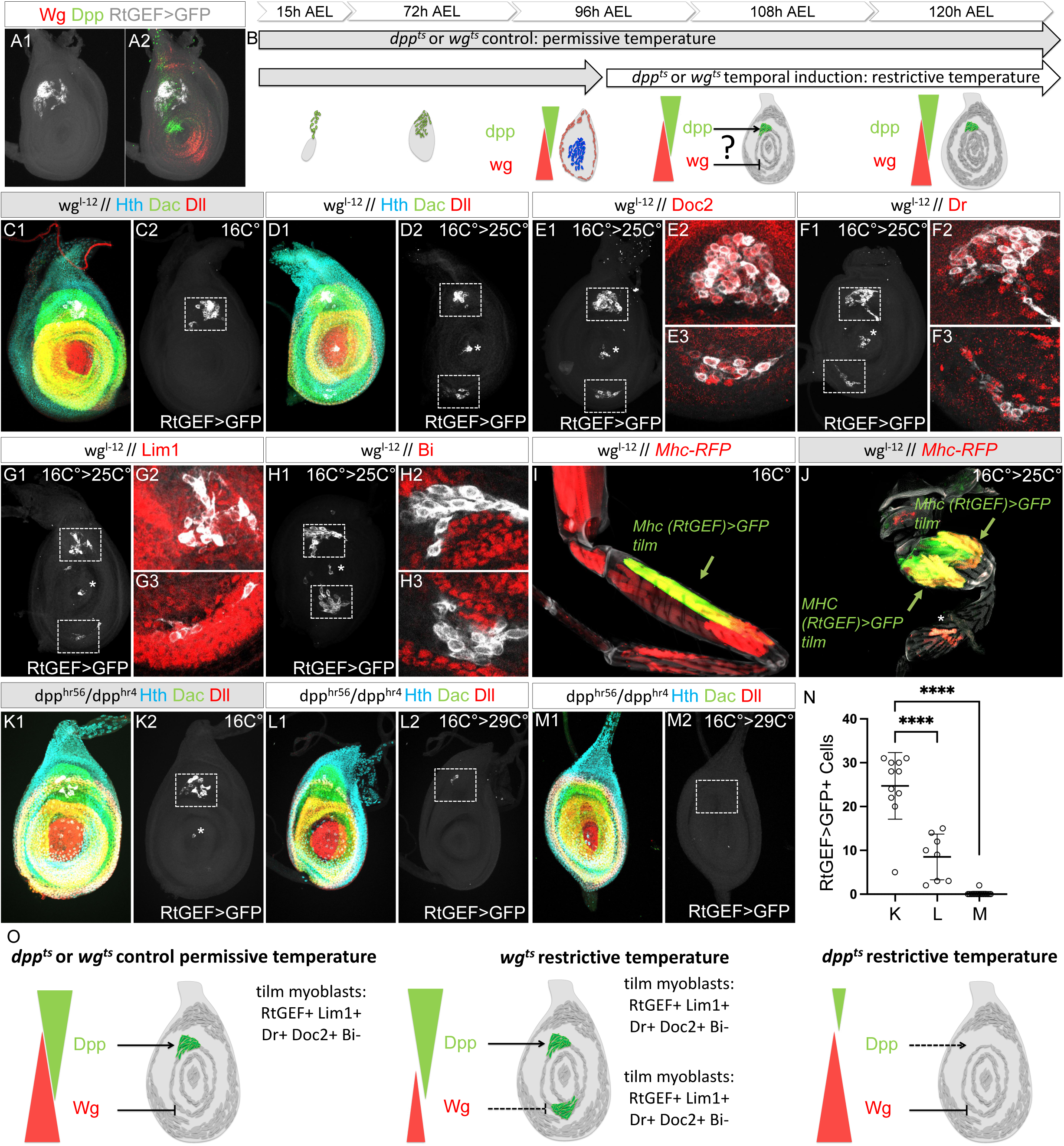
Wg and Dpp act antagonistically to specify MPs to give rise to the tilm, dorsal muscle: **(A1–A2)** Late third instar T1 leg disc genetically labeled with *RtGEF-Gal4>UAS-mCD8::GFP* (gray), immunostained for Dpp (green) and Wg (red). **(M)** Schematic of the conditional *dpp* and *wg* loss-of-function strategy. **(C1–H3)** Third instar leg discs of *wg^1-12^* // mutants expressing *RtGEF-Gal4>UAS-mCD8::GFP* (gray). Discs were immunostained for Dac (green), Dll (red) and Hth (cyan) **(C1-D2,)** or Doc2 (red) **(E1-E3)** or Dr (red) **(F1-F3)** or Lim1 (red) **(G1-G3)** or Bi **(H1-H3).** Samples were maintained at the permissive temperature **(C1-C2)** or shifted to the non-permissive temperature after the specification of central vs peripheral MPs **(D1-H3)**. Note: asterisks show *RtGEF-Gal4>UAS-mCD8::GFP* that we occasionally see in the center of the discs. **(I-J)** Adult T1 leg disc of *wg^1-12^* // mutants labeled with *Mhc-RFP* (red), showing GFP expression (green) in the tilm muscle (arrow) following lineage tracing from *RtGEF-Gal4.* Samples were maintained at the permissive temperature **(I)** or shifted to the non-permissive temperature after the specification of central vs peripheral MPs **(J)**. Note: we could also detect an GFP positive muscle in the tibia some time duplicated when flies where maintained at the restrictive permissive temperature. in contrast when flies were raised at 16^°^C **(I)** this GFP^+^ muscle in WT flies raised **(See** Fig. 4 **T2)** **(K1-M2)** Third instar leg discs of *dpp^hr56^/dpp^hr4^* mutants expressing *RtGEF-Gal4>UAS-mCD8::GFP* (gray). Discs were immunostained for Dac (green), Dll (red) and Hth (cyan). Samples were maintained at the permissive temperature **(K1-K2)** or shifted to the non-permissive temperature after the specification of central vs peripheral MPs **(L1-M2)** at late **(L1-L2)** or early **(M1-M2)** time points. Note 1: asterisks show *RtGEF-Gal4>UAS-mCD8::GFP* that we occasionally see in the center of the discs. **(N)** Graph showing the quantification of the number of GFP^+^ cells in **(K1-M2)**. **(O)** Schematic summary of the phenotypic outcomes observed in panels **(C1-M2).**

**Table 1:**

| Muscle name |  | Phenotypes observed in *bi* KD condition |  |  |  |
| --- | --- | --- | --- | --- | --- |
| Tibia | tarm 1 | fiber number reduced (7/16) | Absent (1/16) |  | Normal (8/16) |
|  | tarm 2 | Absent (2/16) |  |  | Normal (14/16) |
|  | tadm | fiber number reduced (15/16) |  |  | Normal (1/16) |
|  | talm | Fibers attach to the wrong tendon (tadt) (11/16) | Fibers Non attached 1/16 | Fiber number reduced (2/16) | Normal (2/16) |
|  | ltm1 |  |  |  |  |
| Femur | tirm |  |  |  |  |
|  | tidm |  |  |  |  |
|  | tilm |  |  |  |  |
|  | ltm2 | rounded (9/16) |  |  | Normal (7/16) |
| Trochanter | ferm |  |  |  |  |
|  | ferdm |  |  |  |  |
| Coxa | trrm |  |  |  |  |
|  | trdm |  |  |  |  |
|  | trlm |  |  |  |  |

Upon *Lim1* knockdown (KD), tibial muscles are generally weakly affected, though in some cases small fibers are observed targeting incorrect tendons **(Fig. 5G3)**. The most frequent phenotype in the tibia is an increased number of *tarm1* fibers (N=4/10), occasionally resulting in a complete duplication of this muscle **(Fig S6)**. Several femurs appear folded at the level of the *tilm* (N=6/10). In these folded legs, we often observe rounded *tilm* fibers that fail to attach to tendons **(Fig. 5G1, G5-G7)**. This phenotype is likely underestimated, as many animals die in the pupal case late during development, often the posterior part of the body stuck in the pupal case. All examined dead animals showed a folded femur phenotype.

By following TFs expressed in central myoblasts during development, we observed that the expression of TF codes changes dynamically in space and time, giving rise to three distinct populations of myoblasts and one population of neural lamella cells, each characterized by specific TFs signatures **(Fig. 5H)**. For example, myoblasts destined to form the *tilm* muscle express a unique TF code, including *Doc1/2* and *Dr*, both of which show de novo expression at late larval stages. We confirmed the presence of Lim1, Dr, and Doc2 proteins by immunostaining **(Fig. 5I1-K)**. Strikingly, when *Dr* is knocked down in MPs and muscles, only the *tilm* muscle is affected, which becomes atrophic **(Fig. 5 L1-M)**. Most legs analyzed under these conditions were folded.

These results demonstrate that central MPs rely on dynamic and spatially restricted combinations of TFs to define specific distal muscle identities. Individual TFs within these codes, such as *bi*, *Lim1*, and *Dr*, have highly specific and instructive roles, revealing a modular logic underlying muscle diversification in the developing leg.

### Wg and Dpp secreted by the epithelium drive the first step of myoblast specification

Our data reveal that MPs progressively transition from a naïve state through a finely tuned, multistep process that ultimately specifies MPs to generate muscle diversity. The first step of myoblast specification distinguishes MPs located at the center of the disc from those at the periphery, which give rise to distal and proximal muscles, respectively.

Wg (Wnt1) and Dpp (BMP) are two morphogens secreted by the ventral and dorsal epithelium, respectively^24^. We found that MPs at the center of the discs specified to produce telopodites muscles localize directly beneath the sources of both morphogens **(Fig. 6A1-A2)**. Although myoblasts do not secrete these morphogenes, they express the necessary receptors to respond to them **(Fig. S7)**.

To test the role of Wg and Dpp in this process, we used temperature-sensitive alleles of *wg* and *dpp* to inactivate their functions before the initial step of MPs specification (proximal *vs*. distal fate specification). When either Wg or Dpp signaling was disrupted, central myoblasts (Tnc⁺, Unpg⁻, Hth⁻) lost their identity and adopted a peripheral fate (Tnc⁻, Unpg⁺, Hth⁺) **(Fig. 6C1-M)**. In the absence of Wg, the discs were smaller, likely due to both impaired cell proliferation -a well-established function of Wg- and a reduction of the telopodite territory in the epithelium, reflecting Wg’s role in patterning. Nevertheless, the epithelium still expressed Dll, and the central cluster of myoblasts remained localized below this Dll+ Cells, allowing us to identify them as telopodite myoblasts converted into peripheral identity.

These findings indicate that the synergistic action of Wg and Dpp is essential for the initial specification of myoblasts into proximal *vs* distal fates, which appear to represent the ground state **(Fig. 6M)**.

### Wg and Dpp also control the second step of myoblast specification

Following the initial segregation into proximal and distal populations, myoblasts undergo further specification into distinct muscles **(Fig. 5)**. To investigate the developmental logic underlying muscle diversity, we focused on the specification of lineage 11 → 6, which gives rise to the *tilm* muscle.

Myoblasts in cluster 6 express a specific combination of TFs **(Fig. 5)** and localize near the Dpp source **(Fig. 7A1-A2)**, suggesting that Dpp promotes the specification of *tilm* myoblasts. Conversely, Wg may repress their specification in the ventral region.

To test this hypothesis, we inactivated Wg or Dpp signaling after the initial peripheral/central specification step (96 hours AEL). In *dpp* mutants, central MPs failed to adopt the *tilm* fate (cluster 6), as revealed by a strong reduction or absence of RtGEF>GFP⁺ cells depending of the Dpp timing inactivation **(Fig. 7K-N)**. These results indicate that the number of myoblasts specified to form the *tilm* muscle depends on the duration of their exposure to Dpp signaling.

Conversely, in *wg* mutants, we observed a *de novo* cluster of RtGEF>GFP⁺ myoblasts in the ventral region of the disc **(Fig. 7D-H)**. Co-staining with antibodies against TFs expressed in *tilm* myoblasts or other myoblast populations revealed that this ectopic cluster represents a duplication of the *tilm* myoblast identity (Doc^+^, Dr^+^, Lim^+^, bi^−^) in the presumptive ventral domain.

To determine whether this duplication leads to the formation of an additional *tilm* muscle, we lineage-traced both clusters through to the adult stage and observed the presence of a duplicated *tilm* muscle in adults **(Fig. 7I-J)**.

These findings demonstrate that, beyond their role in early myoblast segregation, Wg and Dpp signaling continue to control the second step of myoblast specification by spatially restricting the identity and positioning of *tilm* progenitors, thereby ensuring the correct patterning of adult leg muscles.

## DISCUSSION

### Myoblast are specified before fusion to generate muscle diversity

How individual muscle identities are specified during development remains a central question in developmental biology. In this study, by integrating genetic lineage tracing, single-cell RNA sequencing, and trajectory analysis, we demonstrate that MPs undergo stepwise transitions from a naïve state to acquire distinct identities, thereby specifying muscle diversity prior to fusion. This mechanism shows that MPs and myoblasts are not passive actors simply following the prepattern established by connective tissue cells such as tendons. Instead, disrupting the expression of transcription factors such as *bi* in specific muscle lineages leads to incorrect tendon connectivity, revealing that communication between the two tissues is both specific and, at least in part, intrinsically encoded within muscle lineage.

Lineage tracing revealed that MPs remain multipotent until mid-larval stages, after which they become progressively restricted through a temporally and spatially regulated program. The first major bifurcation separates central and peripheral MP populations within the leg imaginal disc, which subsequently give rise to distal and proximal muscles, respectively. These territories are defined by distinct TF expression profiles: Lim1, Bi, Ko, and Sox100B in the central region, and Hth and unpg in the peripheral domain.

As development proceeds, these broad populations further diversify into transcriptionally distinct subpopulations, each prefiguring the position of specific muscle groups or individual muscles. Intriguingly, we also identified a lineage of mesodermal cells in the distal leg that gives rise to neural lamella cells, suggesting broader developmental potential than previously appreciated. Accordingly, we propose replacing the term AMPs with MPs to better reflect this expanded identity.

We stopped our analysis at early pupal stages since we identified a lineage already committed to forming a single muscle at this stage, validating our model, and demonstrating how specific transcriptional signatures emerge early. Although our current dataset captures early diversification, we expect that later time points would reveal additional fate bifurcations leading to full muscle patterning.

### The driving force of MP/myoblast specification is the morphogens secreted by the epithelium

Finally, our data demonstrate that morphogens secreted by the epithelium are key drivers of myoblast specification, enabling the spatiotemporal coordination of epithelial and mesenchymal development. By following the developmental trajectory of the *tilm* (a distal leg muscle), we found that morphogens function in multiple, temporally distinct phases. Initially, Wg and Dpp act synergistically to specify central myoblasts toward a distal muscle fate. In the absence of either signal, MPs adopt a proximal identity. At later stages, Dpp promotes tilm specification from dorsal central myoblasts, while Wg represses this fate in the ventral region.

Importantly, our findings reveal that myoblasts integrate Wg and Dpp signaling in a time-dependent manner, adjusting their gene expression accordingly. This highlights a mechanism of spatiotemporal signal integration, through which morphogen exposure is interpreted dynamically to guide muscle identity and patterning.

### Evolution of leg appendages in arthropods

In an evolutionary context, our study supports the classical division of the arthropod limb into coxopodite and telopodite^25^. The prevailing theory suggests that the telopodite -the ‘true leg’-evolved from the coxopodite, which is considered to be part of the thoracic body wall. In *Drosophila*, this theory is supported by numerous studies showing that the molecular signature of the epithelial cells of the coxopodite (comprising the coxa and trochanter) closely resembles that of the trunk epithelium, whereas the epithelial cells of the telopodite display a distinct molecular identity.

In the *Drosophila* leg imaginal disc, the central region exposed to both major morphogens, wg and Dpp, gives rise to the telopodite (Dll⁺ and Dac⁺), while the peripheral region differentiates into the prothorax and coxopodite (Hth⁺). This early organization of the leg epithelium is paralleled by a patterning in the mesoderm. Peripheral mesodermal precursors that generate the coxa and trochanter (coxopodite) share transcriptomic signatures with thoracic mesoderm, whereas telopodite MPs (femur, tibia, tarsus) form a transcriptionally distinct population. This supports the view that the telopodite constitutes a true appendage, whereas the coxopodite represents an extension of the body wall.

This mechanism allows the coordinated development of the epithelium and the mesodermal derivatives (the MPs), while also providing evolutionary plasticity. Since the epithelium is the source of morphogens, changes in its patterning will also reshape the patterning of mesodermal derivatives.

### A potential mechanism conserved in vertebrates

In vertebrate limb development, MPCs migrate from the somites into the limb bud, where local signals-largely derived from a prepattern established by connective tissue cells-determine their final positioning. In this classical model, muscle precursors are viewed as developmentally “naïve”, passively following the prepattern established by the connective tissues.

However, several studies challenge this view and instead support a model of early lineage restriction. For instance, MPCs at early developmental stages exhibit bipotency, retaining the ability to differentiate into both muscle and endothelial cells in response to environmental signals^26^. This plasticity mirrors what is observed in *Drosophila*, where mesodermal precursors can also give rise to non-muscle cell types, such as neural lamella cells.

A further layer of evidence suggests that muscle diversification may not result solely from the passive positioning of myoblasts. Adult muscles also display heterogeneity in their sarcomeric organization. For instance, Myosin Heavy Chain (MHC) proteins, the main components of sarcomeres, exist in several isoforms that are differentially expressed across muscle groups under the control of a muscle-specific super enhancer. This regulation suggests that muscles follow distinct developmental transcriptional programs rather than converging on a single default pathway^27^.

Additional evidence for an active specification into specific muscles comes from the spatiotemporal expression patterns of HOXA11 and HOXA13 in vertebrate limb muscle precursors, which suggest that myoblasts are already pre-specified to contribute to particular muscle lineages. Notably, BMP signaling from epithelial cells has been implicated in regulating Hox gene expression in these cells, reinforcing the idea that positional signals influence MPC identity from early developmental stages^28–31^. In line with this, although not explored in depth here, we observed differential Hox gene expression in early myoblasts of the *Drosophila* leg disc, with Antp expressed in central myoblasts and Scr in peripheral ones. These findings suggest that the spatio-temporal control of MP identities by epithelial cells may represent a conserved mechanism guiding muscle lineage diversification across species.

This logic of early lineage restriction may also have implications for human disease. Several muscle-specific myopathies, including muscular dystrophies that selectively affect certain muscle groups while sparing others^32,33^, may reflect an underlying developmental lineage-based organization of muscles.

Such patterns highlight that muscles are not functionally or molecularly homogeneous, but instead may retain distinct embryonic origins and regulatory programs that influence their disease susceptibility.

## Material & Methods

### KEY RESOURCE TABLE

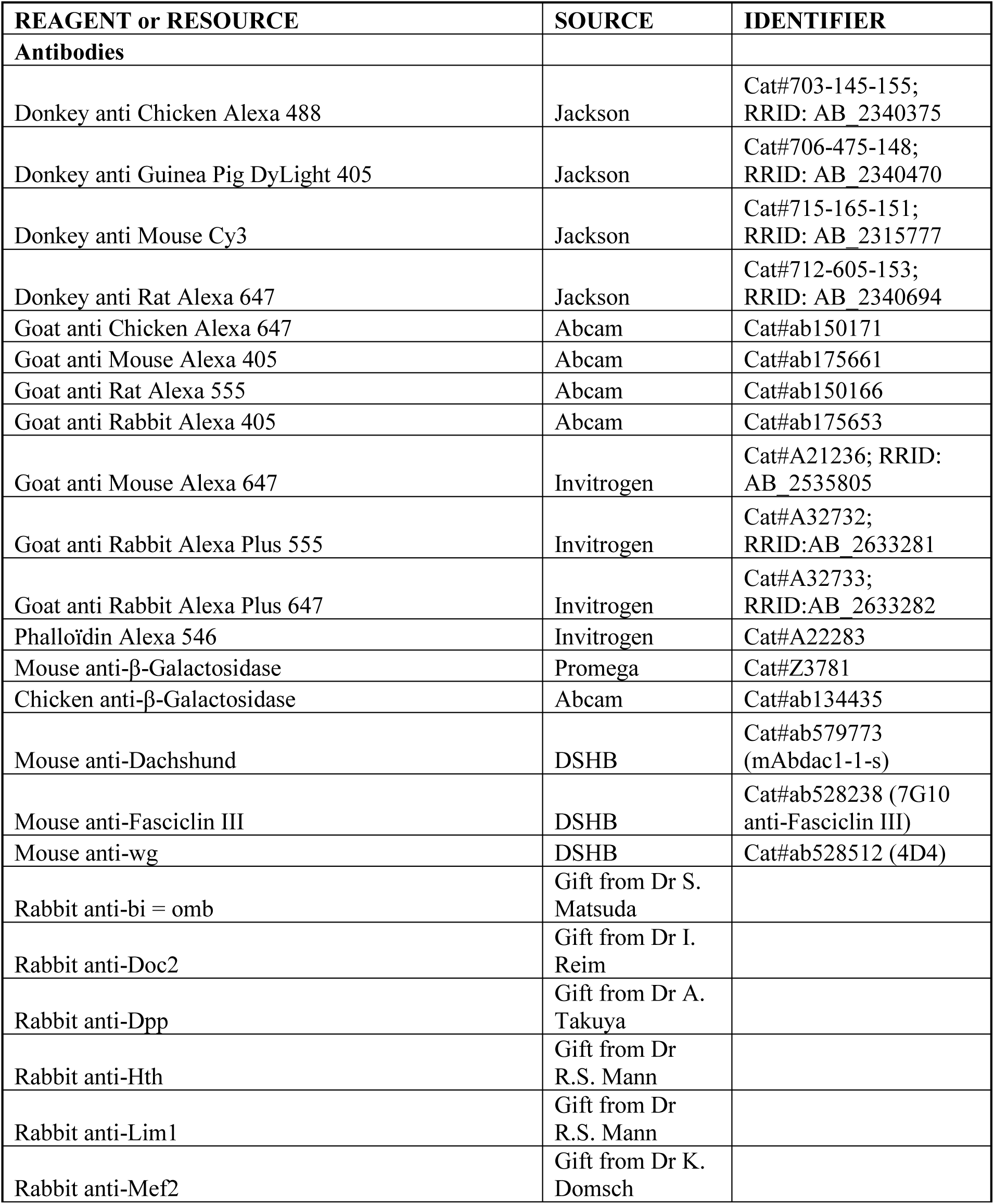
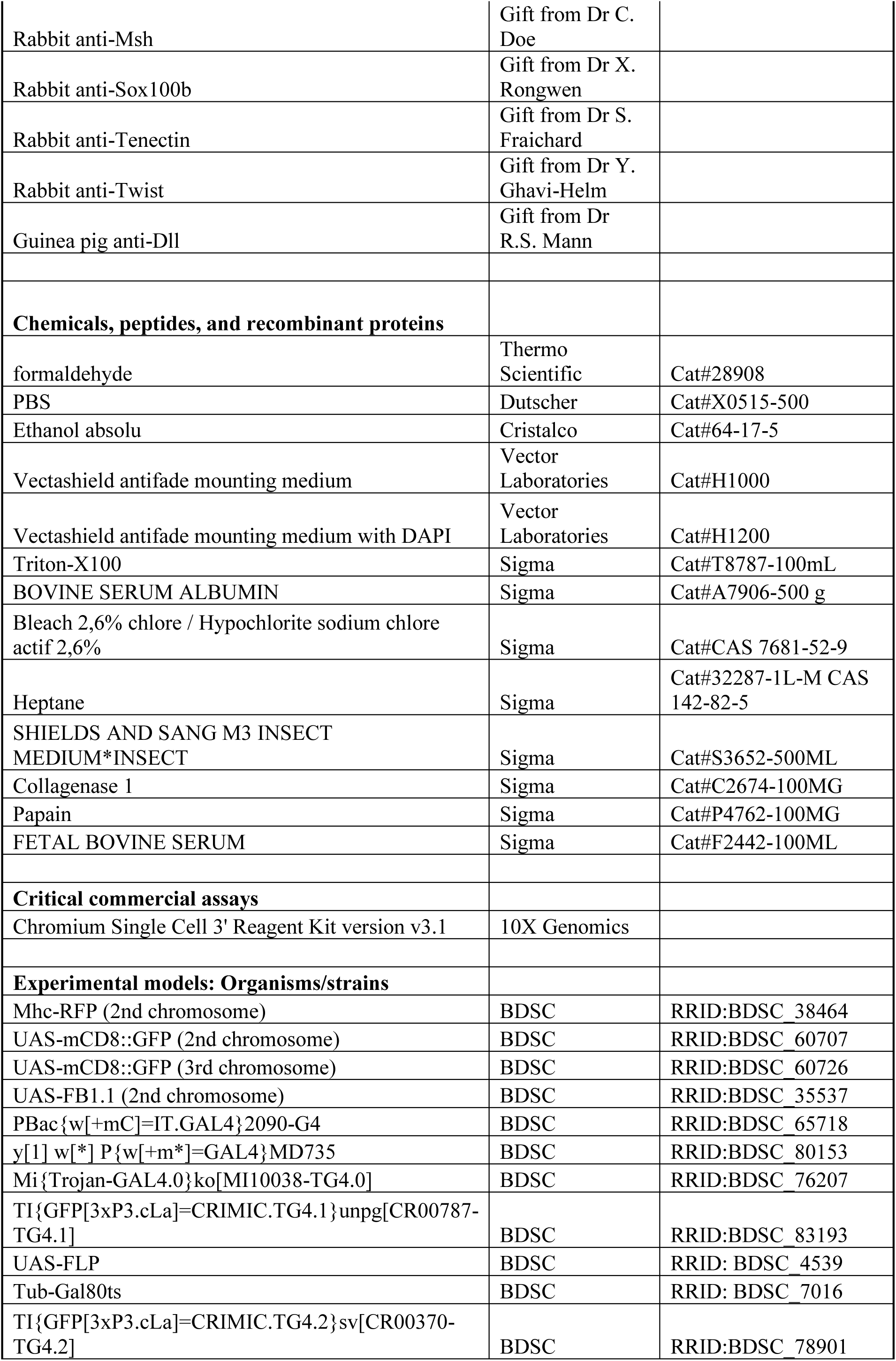
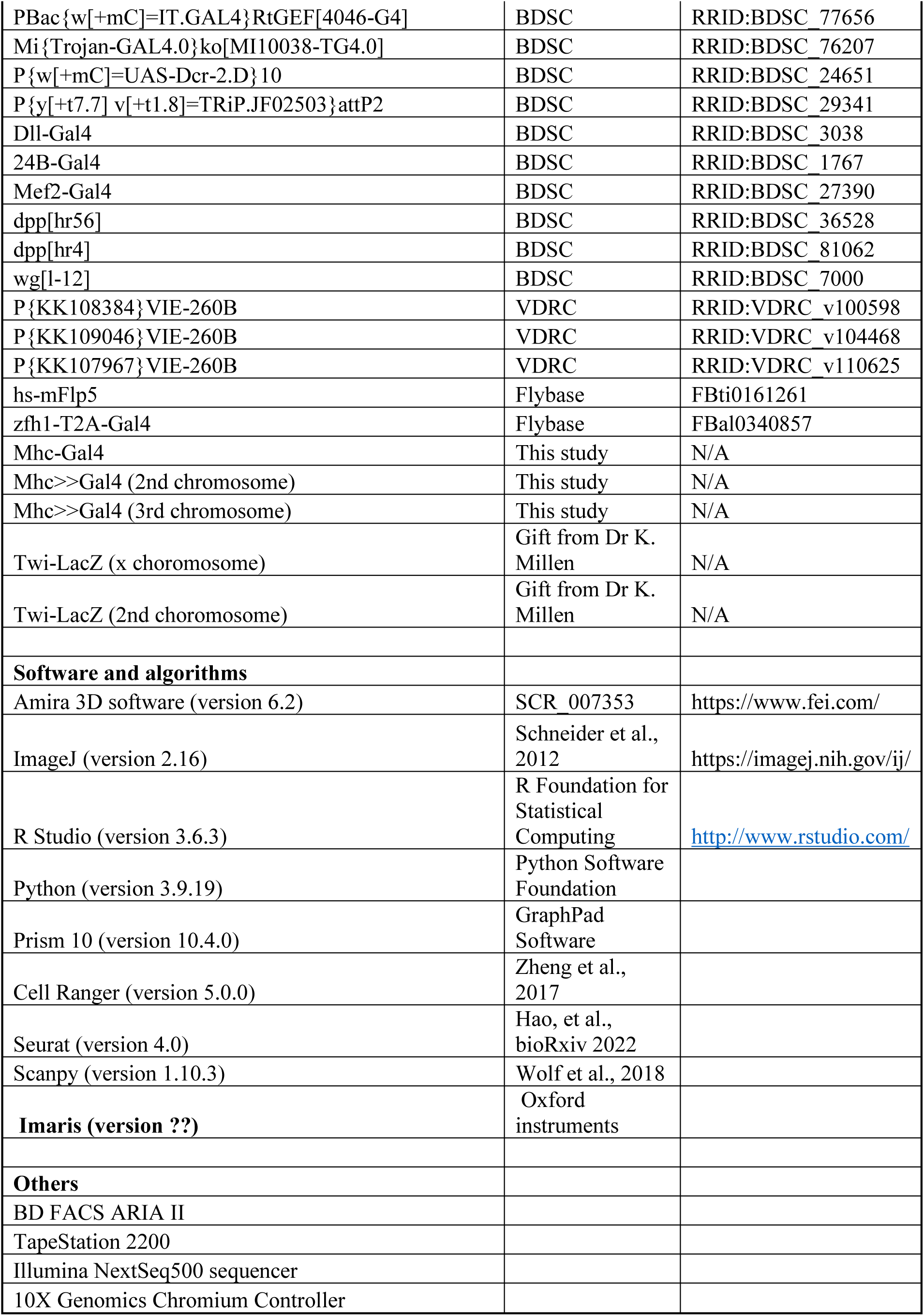

### RESOURCE AVAILABILITY

#### Lead contact

Further information and requests for resources and reagents should be directed to and will be fulfilled by the lead contact, Jonathan Enriquez.

#### Materials availability

All fly stocks made in this study, as well as the nucleotide sequences of the plasmids used, are available from the lead contact without restriction.

#### Data and code availability

DOIs are listed in the key resources table. Any other additional information required to reanalyze the data reported in this paper is available from the lead contact upon request.

### EXPERIMENTAL MODEL AND SUBJECT DETAILS

The experimental model for this study was the vinegar fly *Drosophila melanogaster*. A complete list of strains used is provided in the Key Resources Table and in the Genetic Crosses section for each figure. Unless otherwise specified, flies were maintained on standard cornmeal medium at 25 °C under a 12:12 hours light-dark cycle. Unless stated otherwise, males and females were chosen at random.

### METHOD DETAILS

#### Genetic Crosses for each figure

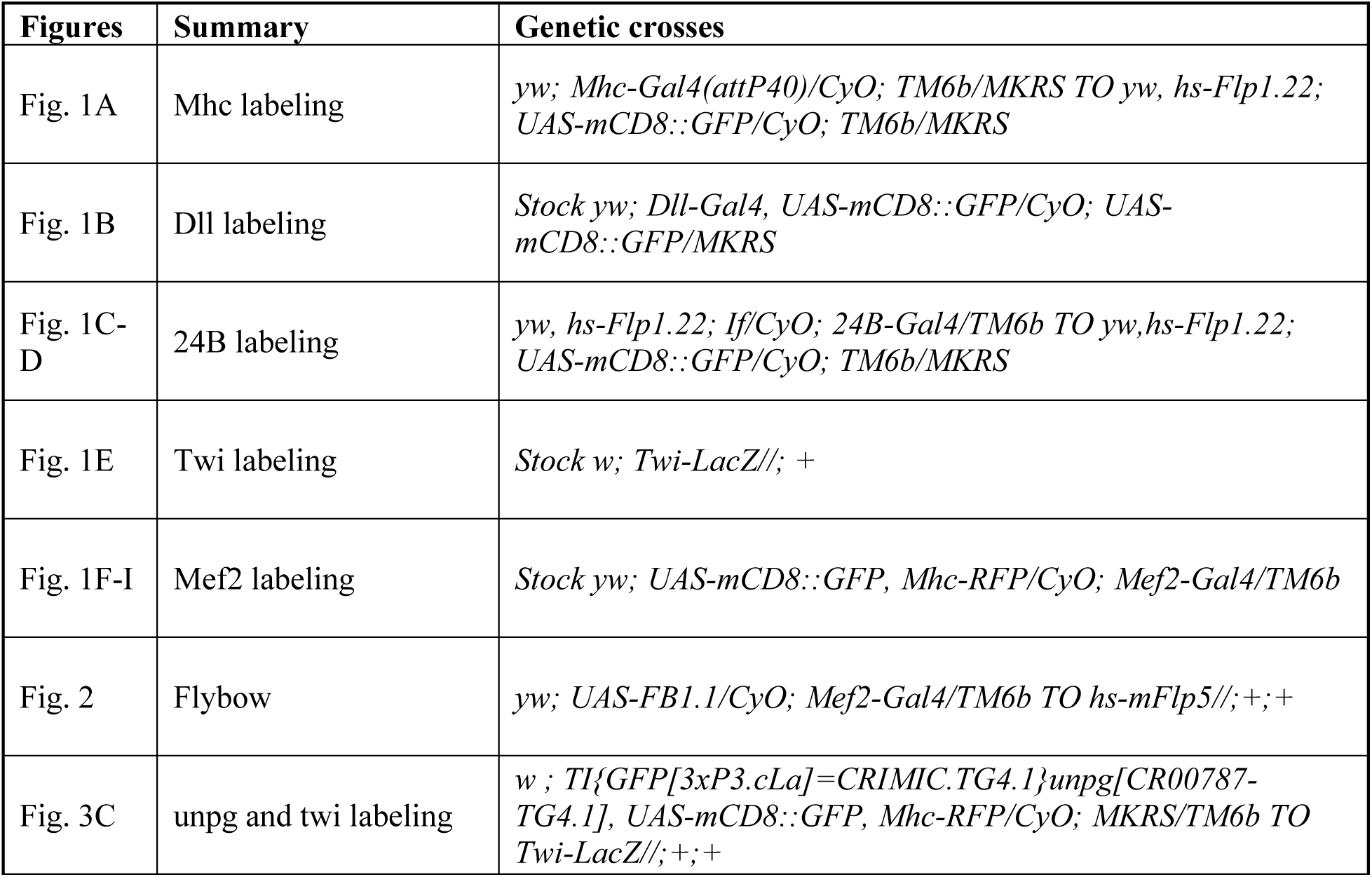
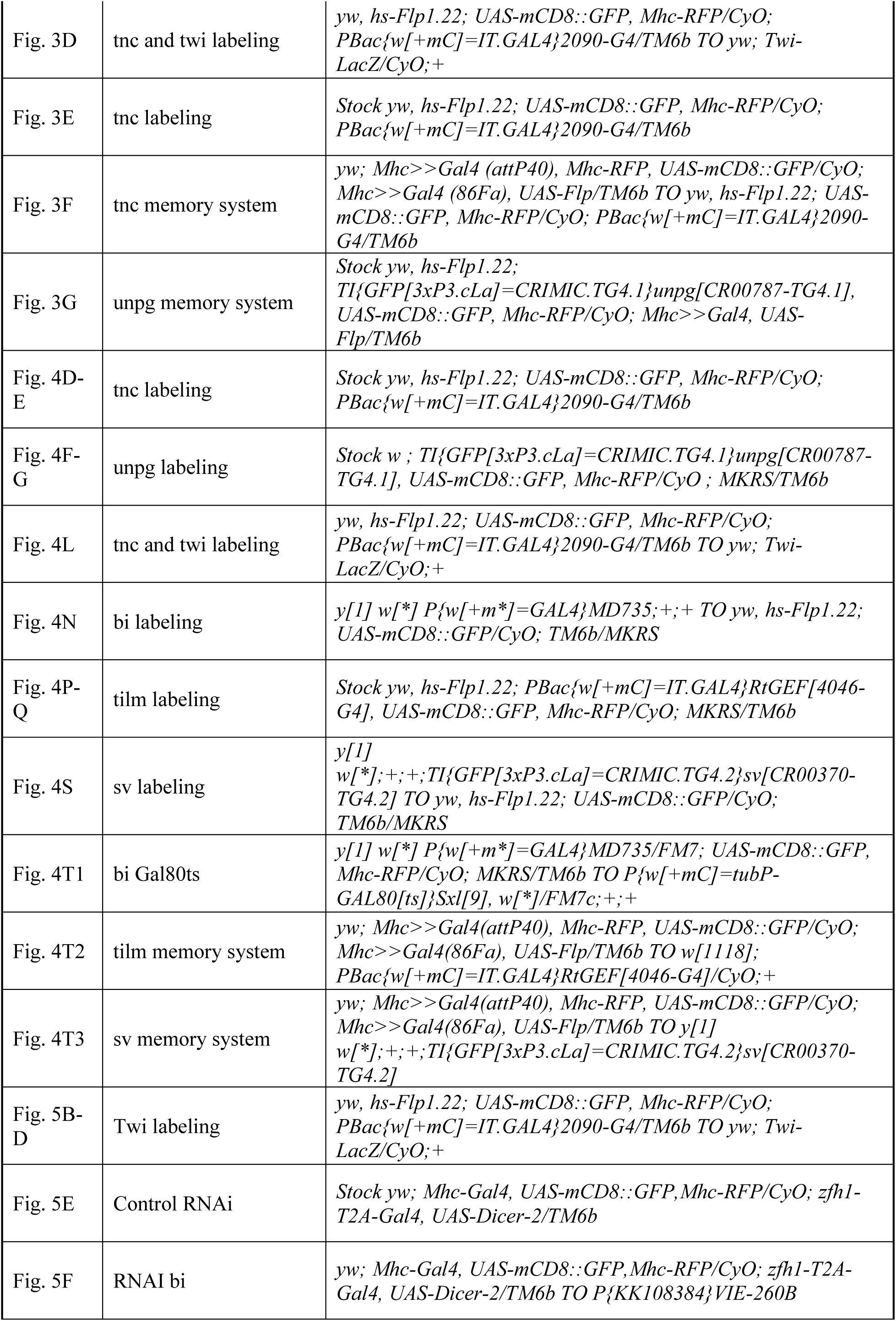
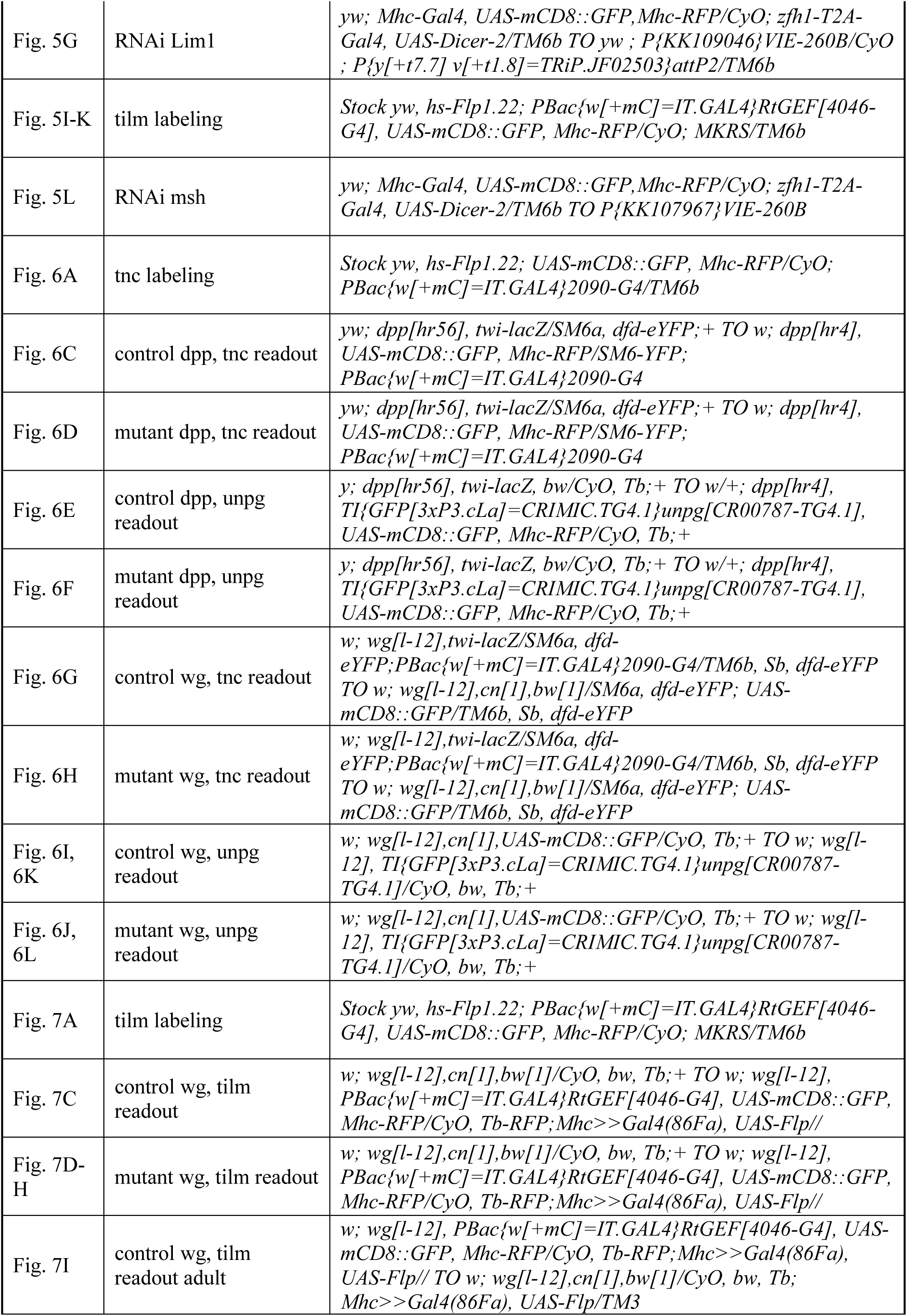
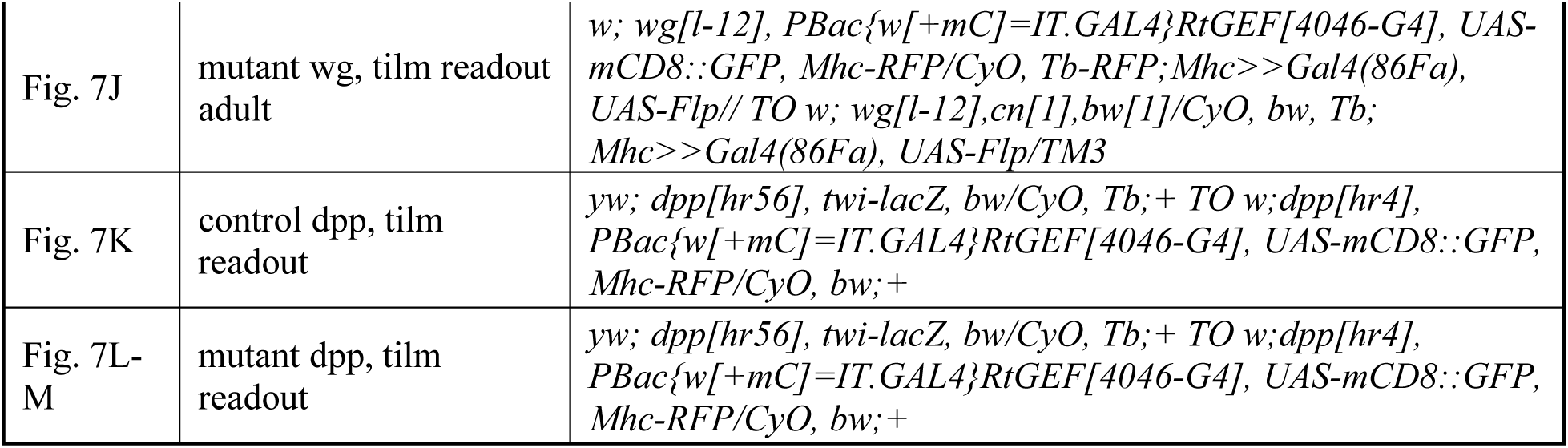

#### RNAi experiments

Crosses were performed at 25 °C, and the progeny were shifted to 29 °C at 24 hours AEL. Adult male and female legs were dissected. Since both sexes displayed similar phenotypes, only T1 legs from males are shown in this study.

#### Leg imaging

Legs were dissected, fixed, and imaged as described in (Guan et al., 2018). We noticed that the internal tendon fluoresces at the same wavelength as the cuticle. To visualize the tendon, the channel corresponding to cuticle autofluorescence was isolated, and all external cuticle signals were manually removed using ImageJ software (version 2.16), leaving only the tendon autofluorescence (pink in **Fig. 5**).

#### Imaging of Prothorax with Attached Forelegs

Heads, abdomens, and mid- and hind-legs were removed in 1X PBS. The thorax, with attached forelegs, was cut along the midline using scissors to obtain two hemi-thoraces. Distal tarsis were removed to improve fixation. Fixation was performed in 4% formaldehyde in 1X PBS for 30 minutes at room temperature. Imaging was performed as described in (Guan et al., 2018).

#### Quantification of the volume of the muscles

The images of Drosophila legs were acquired using a Leica SP8 STED microscope with a 20× objective. After acquisition, the Lightning option in the Leica software was applied to improve the contrast and resolution of fluorescence images. The resulting z-stacks were pre-processed in Fiji (ImageJ) to reduce autofluorescence signal of cuticle in muscle channel, and the volumes of the tibia levator muscle (*tilm*) and femur muscles were quantified using the Surface module in Imaris. At least ten legs per genotype were analyzed.

#### Immunostaining of embryos

Embryos were collected overnight (16 hours) on apple juice agar plates with yeast. They were dechorionated in bleach for 3 minutes, washed with water and then fixed for 45 minutes in a solution of 500 µL of 4% formaldehyde and 500 µL of heptane.

Following fixation, embryos were transferred onto double-sided adhesive tape. The heptane was fully removed, and embryos are left to air dry under a fume hood for 5 minutes to allow strong adhesion.

Next, 0.1% PBST (PBS with 0.1% Triton X-100) was added, and embryos were manually devitellinized using a tungsten needle. A dorsal midline incision was made starting at the posterior end, and gentle pressure was applied to the anterior pole to expelled the embryo.

Finally, embryos were transferred into 1.5 mL of PBST and processed for immunostaining, following a protocol similar to that used for larval or pupal samples.

#### Immunostaining of leg disc during larval and pupal stages

Larvae and pupae were placed in ice-cold 1X PBS in a dissection dish to reduce movement. After a quick rinse with PBS, larvae and young pupae (0 hours After Pupal Formation [APF]) were reverted by cutting the distal region and turning them inside out to expose internal tissues. For pupae at 5 hours APF, cuticle, head and abdomen were first removed, then a dorsal midline incision was made to allow penetration of the fixative. The fat body and gut were removed.

Immediately after dissection, reverted larvae or opened pupae were transferred to ice-cold 4% formaldehyde freshly prepared in PBS.

Once approximately ten reverted larvae or opened pupae were in the cold fixative, the samples were moved to room temperature for fixation: 20 minutes for larvae and 25 minutes for pupae.

Before antibody incubation, samples were washed five times (20 minutes each) with PBST-BSA (PBS with 0.3% Triton X-100 and 1% Bovine Serum Albumin).

After washing, leg discs from larvae or pupae were dissected and incubated in PBST-BSA for 1 hour at room temperature. Samples were then incubated with primary antibodies diluted in PBST-BSA at 4°C overnight with gentle shaking.

Following incubation, samples were washed five times (20 minutes each) with PBST-BSA, then incubated with secondary antibodies (diluted in PBST-BSA) at 4°C overnight with gentle shaking. A final series of five washes (20 minutes each) was performed with PBST.

Finally, samples were mounted on glass slides using Vectashield anti-fade mounting medium (Vector Laboratories), and slides were imaged immediately.

##### Dilution of primary and secondary antibody

All secondary antibodies listed in the resource viability table were diluted at 1:500. Primary antibodies were diluted as follow: Mouse anti-β-Galactosidase (1/500), Chicken anti-β-Galactosidase (1/500), Mouse anti-Dachshund (1/100), Mouse anti-Fasciclin III (1/100), Rabbit anti-bi = omb (1/1000), Rabbit anti-Doc2 (1/1000), Rabbit anti-Dpp (1/100), Mouse anti-en (1/2), Rabbit anti-Hth (1/5000), Mouse anti-lbe (1/500), Rabbit anti-Lim1 (1/50), Rabbit anti-Mef2 (1/750), Rabbit anti-Msh (1/500), Rabbit anti-Sox100B (1/1000, pre-absorbed), Rabbit anti-Tenectin (1/3000), Rabbit anti-Twist (1/500), Mouse anti-wg (1/50), Guinea pig anti-Dll (1/1000), and Phalloïdin Alexa 546 (1/500).

#### Random lineage tracing of MPs using the Flybow system

The Flybow system was used (UAS-Flybow 1.1) to randomly label MPs and trace their lineage through to the adult stage. The UAS-Flybow 1.1 construct enables stochastic expression of fluorescent proteins from a single transgene via excision events mediated by the heat-shock-inducible *hs-FLP5* recombinase. To drive expression of the transgenes, M*ef2-Gal4* was used. In the absence of recombination, the GFP transgene is located adjacent to the UAS sequences in the correct orientation, resulting in uniform GFP expression in all adult muscles. For lineage analysis, mCherry^+^ clones were specifically tracked, which were readily identifiable using a fluorescence stereomicroscope. To ensure that only single recombination events were visualized, experimental conditions were optimized to achieve a low clone induction frequency, with approximately 1.5% of clones observed in the prothoracic segment. Heat shocks were performed at 37°C for 40 minutes at 24 hours after egg laying [AEL], 30 minutes at 48, 72, and 84 hours AEL, and 35 minutes at 96 hours AEL.

#### Dissection of leg disc for scRNA-seq

scRNA-seq were conducted on GFP MPs/myoblasts at different time points during development: 93-96 hours AEL (mid-L3 stage), 108-111 hours AEL, 0 hours APF (White Pupae), 5 hours APF and 12 hours APF.

For each single-cell preparation, a pool of leg discs was collected from the different developmental time points. MPs/myoblasts were labeled with a membrane GFP (*UAS-mCD8::GFP*) under the control of *24B-Gal4* enhancer trap transgene for earlier time points (93-96 hours AEL and 108-111 hours AEL) (*24B* is a transgene expressed mostly in embryonic mesodermal cells) (Brand and Perrimon, 1993; Luo et al., 1994) that is expressed in MPs at early time points, or *mef2-Gal4* transgene for the later points time points (0 hours APF, 5 hours APF and 12 hours APF) (*mef2-Gal4* is a transgene containing the regulatory region of the myocyte enhancer factor 2 during the expression of Gal4). Crosses were performed on day 1, and the next day, an egg collection of 3 hours was made, and the flies were raised at 25°C. Subsequently, T1 leg discs were dissected in less than 1 hour to ensure tissue integrity, in M3 medium. Then, they were transferred, using a glass Pasteur pipette, into a 1.5 ml tube with 450µL of M3 medium.

| Developmental stages | 93-96h AEL | 108-111h AEL | 0h APF | 5h APF | 12h APF |
| --- | --- | --- | --- | --- | --- |
| Number of discs dissected | 48 discs | 96 discs | 82 discs | 80 discs | 80 discs |

#### Cell dissociation before single cell sequencing

The protocol was adapted from Harzer et al.^36^. We added 25 μL of collagenase I and 25 μL of papain to a tube containing M3 medium and the leg discs (ensure the discs have sunk to the bottom of the tube). The tube was then placed in a Thermomixer and incubated for 1 hour at 30°C with shaking at 300 rpm. During the incubation, the contents were gently mixed twice by aspirating 200 μL of the dissociation solution and expelling it forcefully.

After incubation, the dissociation solution was carefully removed without disturbing the discs at the bottom of the tube, ensuring that all discs had sunk. Then, 500 μL of 1X PBS + 5% FBS was added, and the leg discs were mechanically dissociated using a 200 μL pipette tip by pipetting up and down 10–20 times until the solution appeared homogeneous. Finally, the tubes were kept on ice until the FACS step

#### Fluorescence-activated cell sorting (FACS) sorting before single cell sequencing

FACS was performed on a BD FACS Aria II (AniRA-Cytometry platform, SFR Biosciences). Dead cells were excluded by adding DAPI to the sample, and high-quality single cells were sorted into cold 1X PBS supplemented with 5% FBS. Cytometry data were analyzed using BD FACS Diva software version 9.0.1.

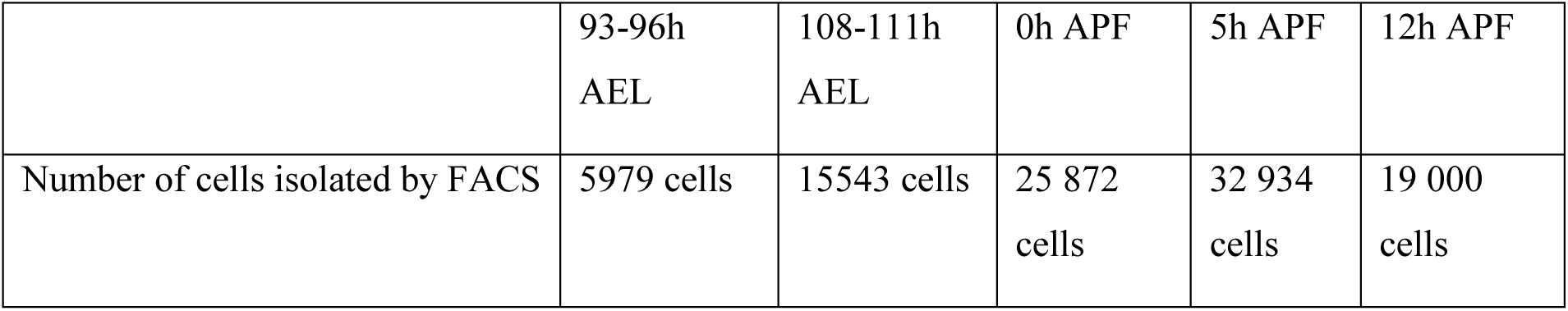

After the isolation of GFP^+^ cells by FACS, the sample is centrifuged for 5min at 500g at 4°C (2017 rpm = 500rcf), and the volume was calculated depending on the number of cells needed for the sequencing experiment. The appropriate supernatant volume was removed, and the pellet was resuspended in the adequate volume. A control of cell concentration using a Malassez chamber was done before the library preparation.

#### Library preparation (10X Genomics protocol)

Preparation of the scRNA library was performed using the 10x Genomics single-cell capturing system. Cells suspensions were loaded onto the 10x Genomics Chromium Controller following the manufacturer’s instructions. Then, single cell cDNA libraries were prepared using the Chromium Single Cell 3’ Reagent Kit version v3.1. For all the time points, a simple index sequence is used, except for time point 2 (108-111 hours AEL) where a dual index is employed. Prior to the sequencing step, the amount, size, and quality of the cDNA were assessed using the TapeStation 2200 (Agilent DNA ScreenTape High Sensitivity System) and the Qubit system.

#### Single-cell sequencing

The resulting libraries were sequenced by the IGFL’s sequencing platform (PSI, Lyon, France) using an Illumina NextSeq500 sequencer (28 bp for R1 and 132 bp for R2).

#### Data pre-processing

Raw sequencing data were generated in base call (BCL) format, which contains data for all libraries within the sequencing run. These BCL files were converted into FASTQ format using the makefastq command from the Cell Ranger software (version 5.0.0; 10x Genomics)^37^. The Cell Ranger pipeline enables demultiplexing of BCL files into FASTQ files corresponding to individual libraries.

Once FASTQ files were generated for each sample, the Cell Ranger pipeline was used for alignment of reads to the reference genome and unique molecular identifier (UMI) counting. Reads were mapped to a customized *Drosophila melanogaster* reference genome (dm6, NCBI RefSeq assembly GCF_000001215.4), modified to include the *UAS-mCD8::GFP* transgene sequence to ensure accurate alignment of reads derived from this construct.

#### Single-cell RNA-seq Data Analysis with Seurat V4

Single-cell RNA-seq data at 93–96 hours AEL were analyzed using the Seurat R package (version 4)^38^, which facilitates quality control, normalization, integration, and exploration of single-cell transcriptomic data. Seurat is specifically designed to identify and interpret sources of cellular heterogeneity and supports integration of diverse single-cell datasets.

Upon loading the data, a Seurat object was created, and initial quality control (QC) metrics were assessed. Cells were filtered based on the following criteria:

- Genes expressed in fewer than 3 cells were excluded (parameter min.cells), to remove uninformative genes.
- Cells with fewer than 200 detected genes were removed (parameter min.features), as these typically represent low-quality cells or empty droplets.
- Cells with total UMI counts >150,000 (or >100 000 for 0 hours APF and 12 hours APF) or <0 were excluded, to remove potential doublets or technical artifacts.
- Cells with >5% mitochondrial gene content were excluded, as high mitochondrial content often indicates stressed or dying cells (mitochondrial genes were identified as those beginning with “MT:”).
- Only GFP^+^ and Twis^+^ cells were retained.

Following cell filtering, data normalization was performed using Seurat’s SCTransform workflow^39^. Highly variable genes, those with significant expression variability across cells, were identified for downstream analysis. A linear transformation (Principal Component Analysis) was applied to reduce dimensionality prior to further processing.

Subsequently, clustering was performed, followed by non-linear dimensionality reduction using UMAP to visualize and explore the data structure. Stochastic neighbour embedding methods aim to project cells into a low-dimensional space while trying to preserve local neighbourhood structures, such that cells within the same cluster are positioned close to one another.

Finally, Seurat was used to identify marker genes defining each cluster through differential expression analysis. By default, Seurat identifies both positive and negative markers for a given cluster compared to all other cells. This analysis can also be extended to compare groups of clusters or perform global comparisons across the dataset.

The code source for the Seurat V4 analysis is available on the following GitLab repository (http://gitbio.ens-lyon.fr/igfl/enriquez/myo_d/clustering_analysis.git). A Docker image is also provided to reproduce the results of the analysis.

#### Single-cell RNA-seq Data Analysis with SCANPY

In parallel with the Seurat workflow, an alternative analysis strategy was implemented using the SCANPY toolkit^40^ to integrate all samples and perform downstream single-cell RNA-seq analysis.

Datasets from multiple time points (108-111 hours AEL, 93-96 hours AEL, 0 hours APF, 5 hours APF, and 12 hours APF) were combined into a single integrated dataset for joint analysis.

The initial preprocessing steps involved filtering out low-quality cells and uninformative genes Cells with mitochondrial gene percentages exceeding 20% were excluded to remove low-quality or dying cells. Additional filtering criteria removed cells with fewer than 600 or more than 5000 detected genes, and cells with total counts above 150000. Genes expressed in fewer than 3 cells were also removed.

For specific developmental stages, targeted filtering was applied:

- At 93–96 hours AEL hours AEL and 108-111 hours AEL, only cells expressing at least one of **GFP** or **Twi** were kept.
- At 0 hours APF, 5 hours APF, and 12 hours APF, only cells expressing at least one of the following genes: **GFP**, **Mef2**, or **Twi**, were retained.

Filtered data were normalized by scaling each cell to a total of 10,000 counts. The normalized counts were then log-transformed (log(x + 1)) to stabilize variance for downstream analysis. Finally, gene expression values were scaled to unit variance and zero mean.

Batch effects across samples were corrected using the BBKNN algorithm^41^, which preserves the local structure of the data while adjusting for technical variability between batches.

A first analysis and clustering were performed on each time point individually, and differential gene expression analysis was conducted similarly to the Seurat-based workflow to identify marker genes associated with each cluster.

Further analysis was conducted on the combined dataset withs all the time points.

Further details and the source code for the SCANPY analysis is available on the following Git repository: https://github.com/spatial-cell-id/Guillermin_et_al.git

### Single-cell RNA-seq Pseudotime Analysis with PAGA

To investigate the temporal evolution of cellular states and connect different developmental time points, pseudotime analysis was performed using the Partition-based Graph Abstraction (PAGA) algorithm implemented in Scanpy^22^. PAGA constructed a graph-based representation of cellular trajectories, providing insight into the connectivity and progression between clusters within a heterogeneous cell population. This approach facilitates the reconstruction of lineage relationships and the temporal ordering of cells across developmental stages.

PAGA analysis was applied to a subset of the data including clusters 4, 6, 11, 13, 14, and 15 (**Fig. 4**), which represent central MPs/myoblasts contributing to the development of telopodite muscles. Cluster 11, corresponding to central MPs at the earliest time point, was selected as the root for pseudotime inference **(see Fig. 3 and Fig. 4A1)**.

PAGA connectivities tree and ForceAtlas2 (FA2) map pseudotime color-coded were generated across all central MP/myoblast clusters. Lineages were defined using the following criteria:

1. The strongest connection between clusters in PAGA connectivities tree graph.
2. The shortest path in the PAGA connectivities tree graph between clusters.
3. Coherent directionality observed in the ForceAtlas2 (FA2) map colored by normalized pseudotime.

To refine lineage directionality, pseudotime was also visualized on individual lineages, allowing us to distinguish trajectories that evolve at different speeds. Of note, cluster 14 was further subclustered, into 14_1 and 14_2, to improve resolution of the lineage leading to tibial and femoral muscles.

Finally, clusters were color-coded on the FA2 map using pseudotime-matched color gradients. This visualization revealed consistent temporal dynamics across lineages, supporting the conclusion that normalized pseudotime accurately reflects the developmental progression of central MPs.

Further details and the source code for the PAGA analysis is available on the following Git repository: https://github.com/spatial-cell-id/Guillermin_et_al.git

#### Temporal induction of wg and Dpp loss of function

Thermosensitive alleles was used for both *dpp* and *wg* experiments. The allele *wg[IL114]*, referred to here as *wg[l-12]*, was described in Van den Heuvel et al.^42^. The thermosensitivity of the heteroallelic combination *dpp[hr4]/dpp[hr56]* was reported by Hsiung^43^, with the molecular nature of *hr56* and *hr4* detailed by Wharton et al.^44^. To facilitate genotype preparation, PCR assays were developed to distinguish both *wg[l-12]* and *dpp[hr4]* alleles, which enabled to sort recombinant chromosomes efficiently. The *wg* mutant chromosome obtained from the Bloomington stock was homozygous lethal; through recombination, the lethal mutation was separated from the *wg* allele. Mutant crosses were maintained at the respective restrictive temperatures, 25°C for *wg* and 29°C for *dpp*, with control groups kept at 16°C. Egg collections were performed in 3-4 hours intervals, and developmental timing was adjusted accordingly to temperature to ensure that embryos and larvae reached the appropriate developmental stages before temperature shifts were applied (**see Fig.6 and 7**). Mutant animals were sorted on the presence of absence of Tb phenotype when using Tb marked balancer chromosome^45^, or with YFP marked balancer chromosome^46^.

*dpp[hr4]/dpp[hr56] or wg[l-12]* samples were shifted to the non-permissive temperature either before or after the specification of central *vs* peripheral MPs.

- Shift before specification:

– Dpp mutant condition: 3 days AEL
– Wg mutant condition: 6-7 days AEL
- Shift after specification:

– Wg mutant condition: 10 days AEL
– Dpp mutant condition: 8 days + 14 hours AEL (early time point) or 9 days AEL (late time point)

#### Image acquisition

For immunostained leg discs, multiple 0.5µm-thick sections in the z axis were imaged with a Leica SP8 confocal microscope using a 40x glycerol objective. Multiple 1.0µm-thick sections in the z axis for adult legs were images with a Leica SP8 or a Zeiss LSM780 confocal microscope, using 20X or 25X glycerol objectives.

A fluorescent stereomicroscope Leica M205 FA was used to select cherry positive clones **(Fig. 2**)

#### Software

Confocal pictures were processed and analyzed with Image J software (version 2.16), Imaris, and 3D reconstruction were performed using Amira 3D software (version 6.2).

Graphs and statistical visualizations were generated using Prism 10 (version 10.4.0).

Single-cell RNA sequencing data were processed with Cell Ranger (version 5.0.0) and analyzed using Seurat (version 4.0, R-based version 4.2.1) and Scanpy (Python-based, version 3.9.19).

#### Schematics

All schematics were done with Microsoft PowerPoint.

#### Cloning

*Pattb-MHC-DSCP-5’FRT-stop-3’FRT-Gal4-hsp70.* The MHC enhancer^47^ was excised using Acc65I (5’) and XmaI (3’) from a Bluescript vector (a gift from Frank Schnorrer) and cloned into a Pattb-Gal4-hsp70 vector (a gift from the Mann lab) in front of the Gal4 sequence, which had been opened with NheI (5’) and SbfI (3’), resulting in the Pattb-MHC-Gal4-hsp70 vector. In a second step, a 5’FRT-stop-3’FRT cassette was excised from another vector (a gift from the Gary Struhl lab) using KpnI and inserted between the MHC enhancer and the Gal4 cassette in the Pattb-MHC-Gal4-hsp70 vector, which had been opened with KpnI. The resulting Pattb-MHC-DSCP-5’FRT-stop-3’FRT-Gal4-hsp70 construct was inserted into chromosomes II and III by injection into embryos carrying the attP40 or attP86F landing sites.

## ACKNOWLEDGMENTS

We thank Dr Carlos Estella and Dr Cedric Soler for comments on the manuscript. This work was supported by the ANR grant DevandMaintain (#ANR-20-CE16-0007) to JE, the SFM (Société Française de Myologie), the EquipEx+ Spatial-Cell-ID under the “Investissements d’avenir” program (#ANR-21-ESRE-00016) to J.E and Y.G.. We acknowledge the contribution of SFR Biosciences facilities (UAR3444/CNRS, US8/INSERM, ENS de Lyon, UCBL): Arthro-tool facility and AniRA-Cytométrie. We the help of the staff of the Plateau Technique AniRA-Cytométrie, SFR Biosciences (US8/UAR3444), especially M. Sébastien Dussurgey, for assistance with the low cytometer. We acknowledge the contributions of the CELPHEDIA Infrastructure (http://www.celphedia.eu/), especially the center AniRA in Lyon. We thank the CBPsmn for providing free access to the computing clusters and the microcopy facility platform of the IGFL.

## Author contributions

Conceptualization, J.E; methodology, J.E; investigation, C.G, V.T, M.B, A.L, D.Z, S.S, L.G, I.S, G.M, E.T, Y.G, B.G, S.H, S.V, J.E; writing original draft, J.E; writing, review & editing, J.E; funding acquisition, J.E, Y.G; resources, J.E; supervision, J.E.

---

**Figure S1:**
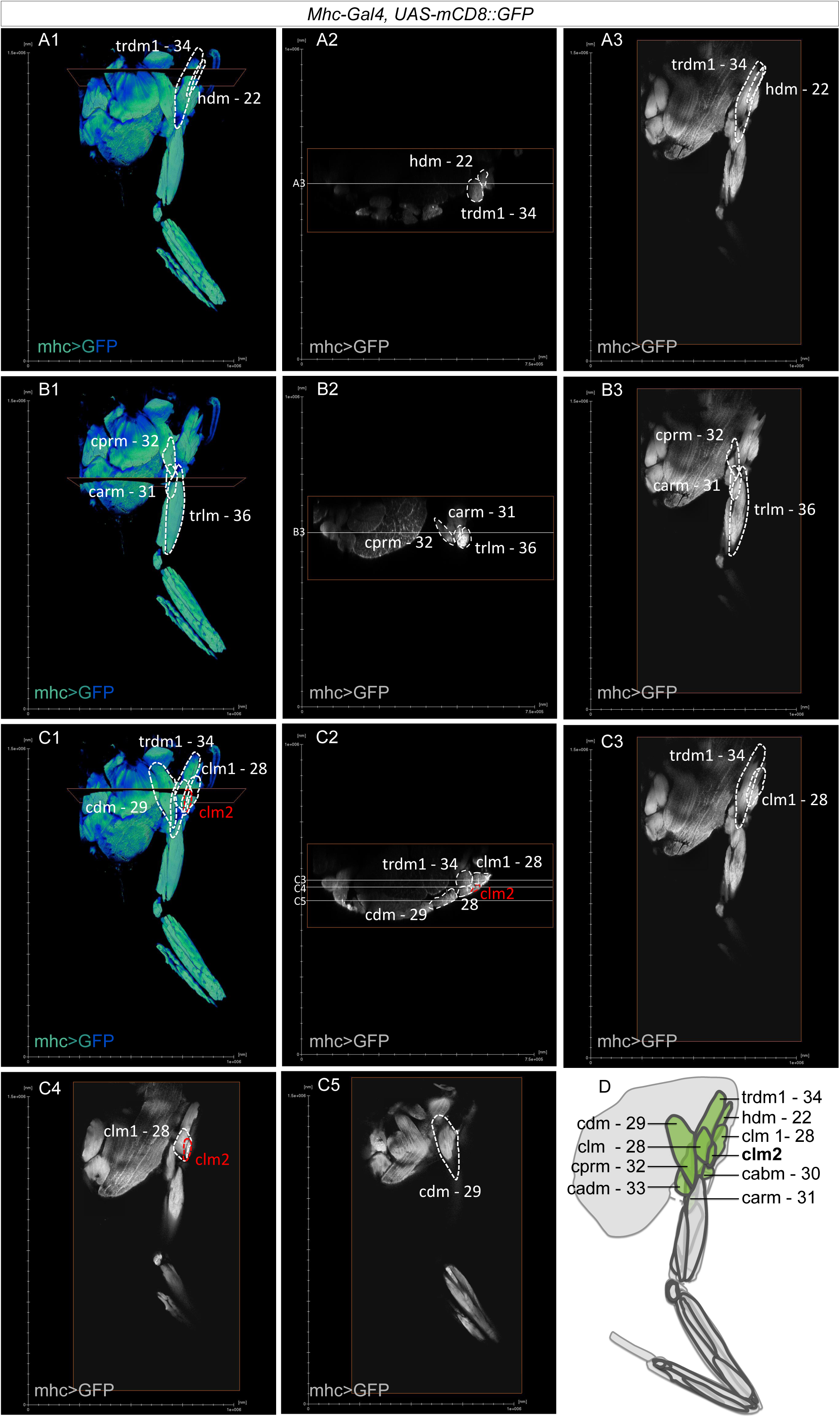
Description of the prothoracic muscles. **(A1, B1, C1)** 3D renderings of confocal sections showing the prothorax and the attached T1 leg, with muscles genetically labeled using *Mhc-Gal4>UAS-mCD8::GFP*. **(A2, B2, C2)** Transverse sections; the positions of these sections are indicated in **(A1, B1, C1)**. These three transverse views were selected to highlight the positions of prothoracic muscles, outlined by dotted lines. A previously undescribed muscle, **clm2**, is shown in red. **(A3, B3, C3, C4, C5)** Sagittal sections; their positions are indicated by white lines in **(A2, B2, C2)**. These sections highlight the positions of the prothoracic muscles outlined with dotted lines. The muscle **clm2**, not described before, is marked in red. **(D)** Schematic representation of all adult thoracic muscles. Prothoracic muscles described in this figure are shown in green. Muscle nomenclature follows Soler and Laurence article^2,3^: muscle; hd: head dorsal; c: coxa; tr: trochanter; fe: femur; ti: tibia; ta: tarsal; l: levator; d: depressor; ab: abductor; ar: anterior rotator; pr: posterior rotator; ad: adductor; r: reductor; lt: long tendon or Miller’s nomenclature^48^: numbers.

**Figure S2.**
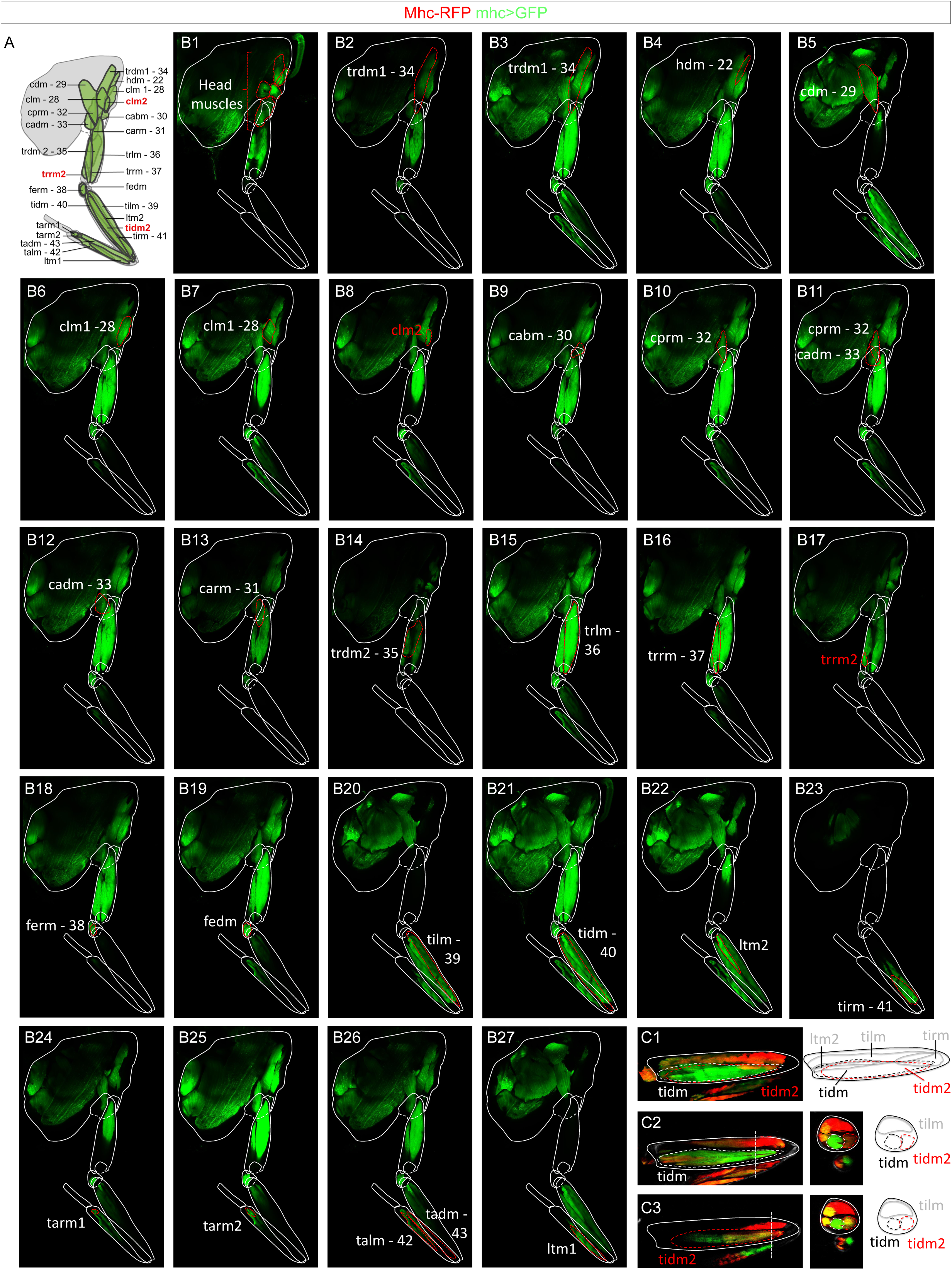
Description of the leg muscles. **(A)** Schematic representation of all adult thoracic T1 leg muscles. Muscle nomenclature follows Soler and Laurence article ^2,3^: muscle; hd: head dorsal; c: coxa; tr: trochanter; fe: femur; ti: tibia; ta: tarsal; l: levator; d: depressor; ab: abductor; ar: anterior rotator; pr: posterior rotator; ad: adductor; r: reductor; lt: long tendon or Miller’s nomenclature^48^: numbers. **(B1-B27)** confocal sections showing the prothorax and the attached T1 leg, with muscles genetically labeled using *Mhc-Gal4>UAS-mCD8::GFP*. **(C1-C3)** Adult femur of T1 legs genetically labeled with *MHC-RFP* (red) and *UAS-mCD8::GFP* (green) under the control of *MHC-Gal4*. **(C1)** shows a maximum projection; **(C2–C3)** are two confocal sections through the tidm or the deeper tidm2. To the left of **(C1)** is a schematic of the femoral muscles. To the left of **(C2–C3)** are two different cross-sections with their corresponding schematics. The positions of the cross-sections are indicated in **(C2–C3)**.

**Figure S3.**
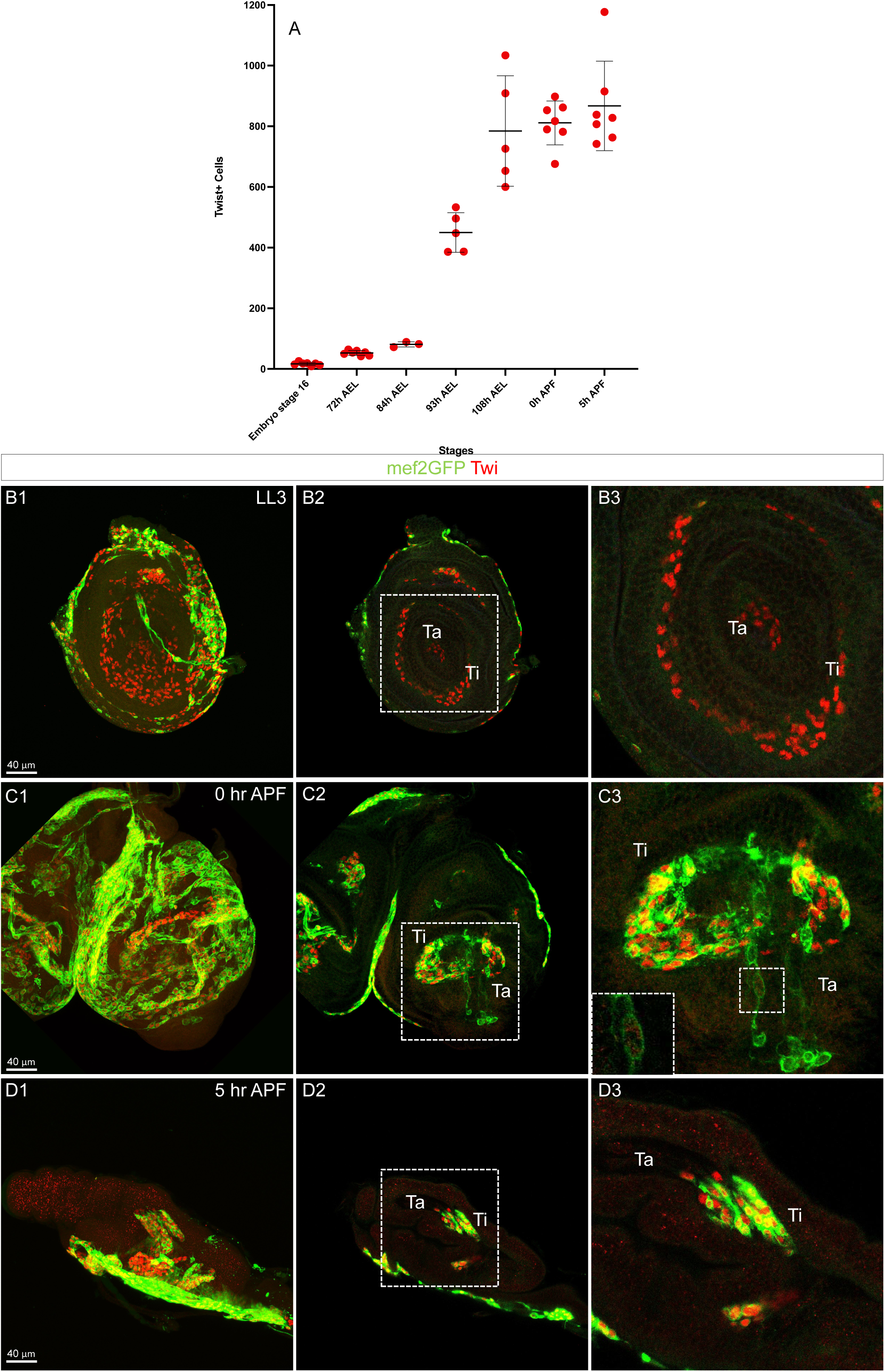
Dynamic of *Mef2* and Twist expression during late larval stages and early pupal stages. **(A)** Graph of the number of Twist^+^ cells during development. **(B1-D3)** T1 leg discs immunostained for Twist (red) and genetically labeled with GFP (green) under the control of *Mef2-Gal4*. **(B1, C1, D1)** are maximum projection, **(B2, D2)** are confocal sections, **(C2)** is a maximum projection of few confocal sections. **(B3, C3, D3)** show enlargement of the dotted box in **(B2, C2, D2). (B3)** shows Twist^+^ Mef2^−^ cells in the tarsus, **(C3)** shows a weak Twist^+^ cells and weak Mef2>GFP^+^ cells in the tarsus, **(D3)** shows the absence of Twist^+^ and Mef2>GFP^+^ cells in the tarsus, **note** depending on the sample weak Mef2>GFP^+^ can be detected as shown in **(Fig. 1, G2).**

**Figure S4.**
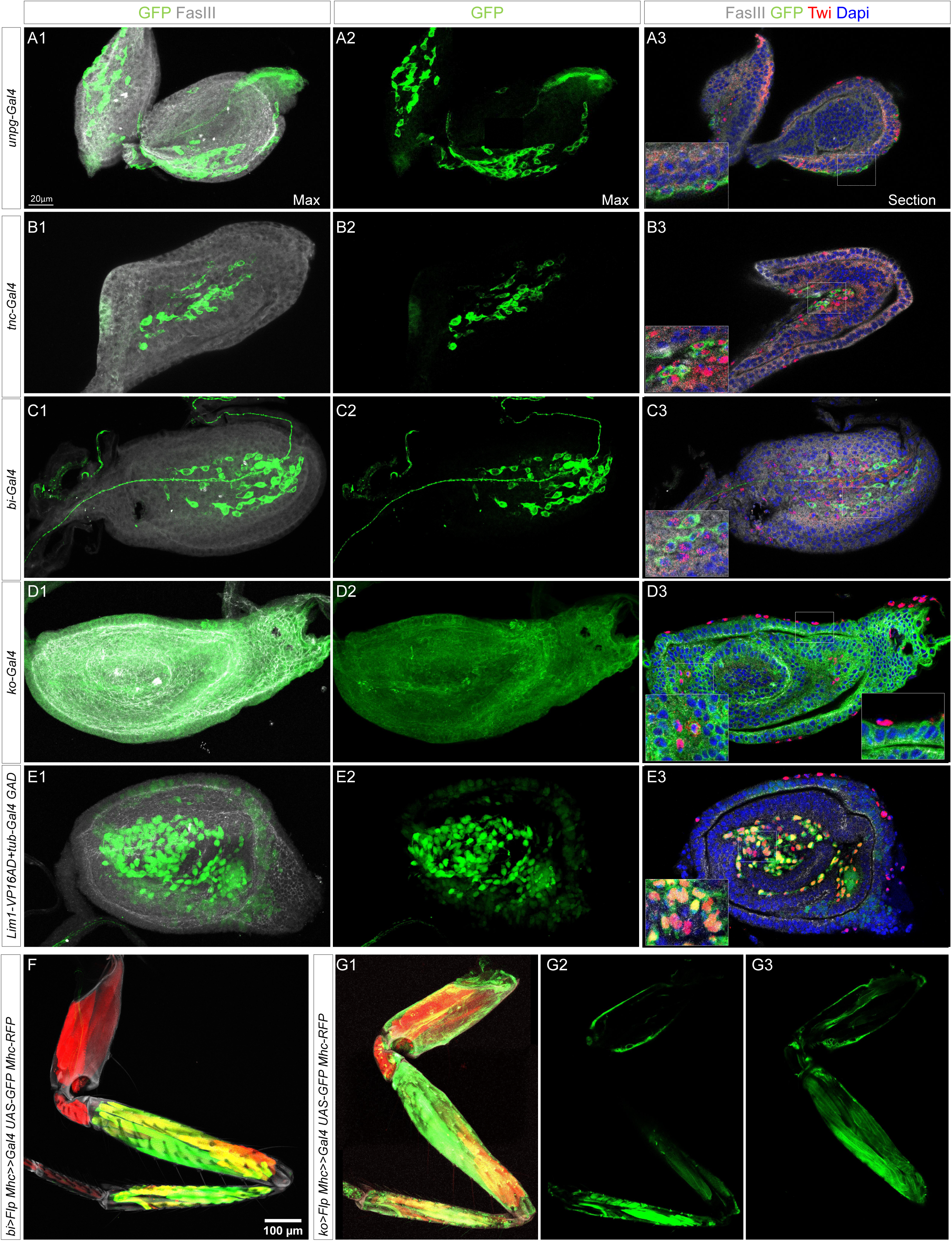
Recovering the spatial information and determining the identity of the central and peripheral MP clusters. **(A1-E3)** T1 leg discs at 96 hours AEL immunostained for FasIII (grey) and Twist (red), counterstained with DAPI (blue), and genetically labeled with GFP (green) under the control of *unpg-Gal4* **(A1-A3),** *tnc-Gal4* **(B1-B3),** *bi-Gal4* **(C1–C3),** *ko-Gal4* **(D1–D3),** and *Lim1-VP16AD + tub-Gal4DBD***(E1-E3).** **(A1–E2)** show maximum projections; **(A3-E3)** are corresponding confocal sections. The bottom left dotted boxes in **(A3, B3, C3, D3, E3)** highlight GFP^+^ and Twi^+^ MPs; the bottom right box in **(D3)** highlights GFP^−^ but Twi^+^ MPs showing the absence of Ko in peripheral MPs. **(F1-G3)** T1 legs labeled with *Mhc-RFP* (red) showing GFP expression in distal and proximal muscles following lineage tracing activated in MPs using *bi-Gal4* **(F)** or *ko-Gal4* **(G1-G3),** respectively. (**F–G1)** are maximum projections; **(G2, G3)** are confocal sections.

**Figure S5.**
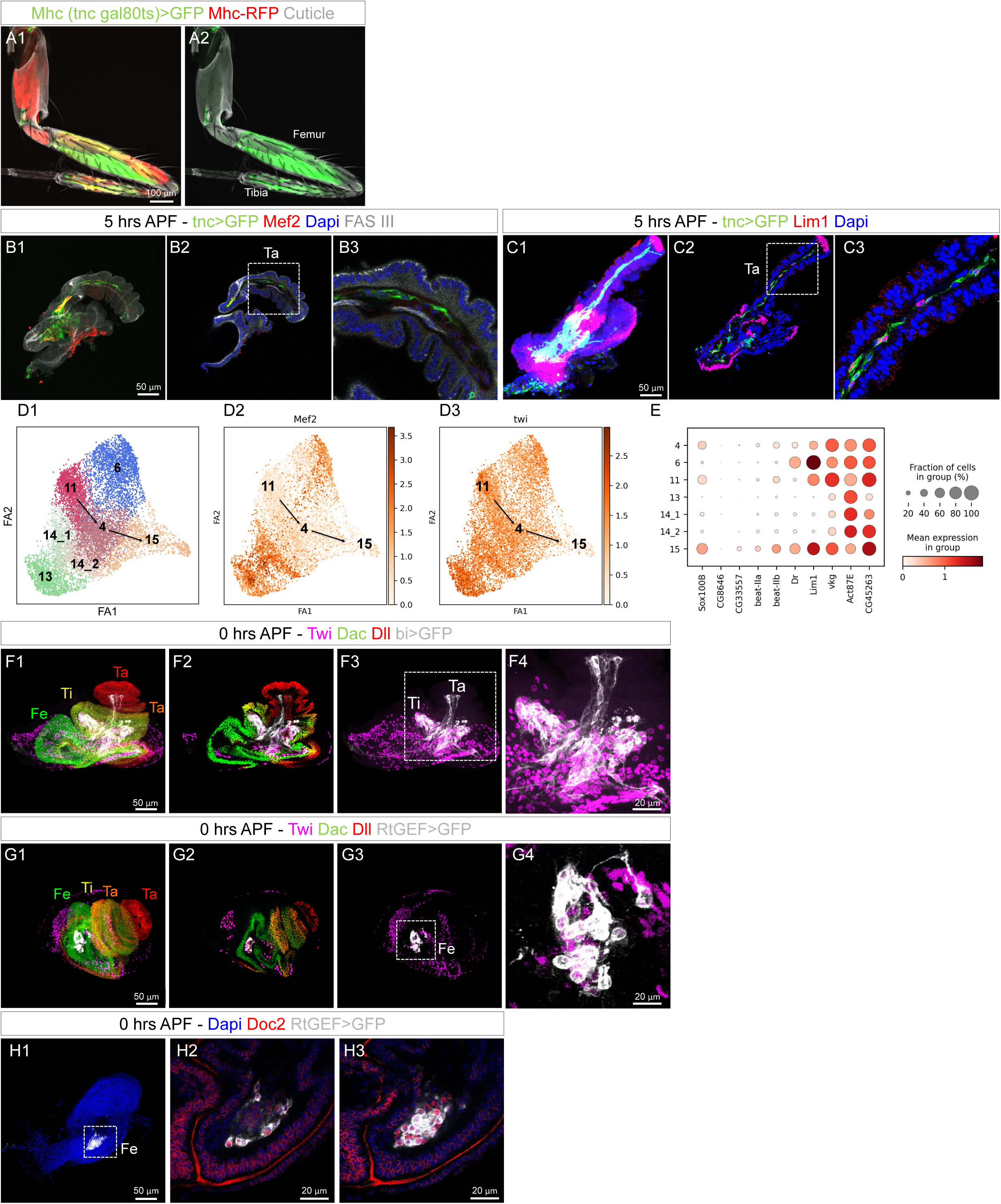
Recovering the spatial information and determining the identity of MP clusters at pupal stages. **(A1–A2)** T1 legs labeled with *Mhc-RFP* (red), showing GFP expression in distal and proximal muscles following lineage tracing activated in MPs using *tnc-Gal4*. *tubP-GAL80^ts^* was used to activate lineage tracing only after 0 hours APF. Note: This late induction is sufficient to label all telopodite muscles, confirming that *tnc-Gal4* is a robust marker for telopodite lineages even at later stages. **(B1-B3)** Immature T1 legs at 5 hours APF immunostained for Fas III (grey) and Mef2 (red), counterstained with DAPI (blue). **(B1)** is a maximum projection; **(B2)** is a confocal section; the dotted box in **(B2)** is enlarged in (**B3)**. Note: tnc>GFP^+^ Mef2^−^ cells are present in the tarsus **(B3)**, and also in the immature femur and tibia (not shown). **(C1-C3)** Immature T1 legs at 5 hours APF immunostained for Lim1 (red), counterstained with DAPI (blue). **(C1)** is a maximum projection; **(C2)** is a confocal section; the dotted box in **(C2)** is enlarged in **(C3)**. Note: tnc>GFP^+^ Lim1^+^ cells are present in the tarsus **(C3)**. **(D1-D3)** FA2 map showing distinct central clusters. **(D1)** Arrows indicate the 11→4→15 lineage; **(D2-D3)** Expression patterns of Mef2 and Twi, respectively. **(E)** Dot plot showing the expression of neuronal lamella markers. Note: The absence of Mef2 (**B1–B2, D2)**, inhibition of Twi **(D3 and see** Fig. 4), and the expression of neuronal lamella markers (**E, C1-C3, and** Fig. 4) indicate that the 11→4→15 lineage gives rise to the neuronal lamella. **(F1-F4)** Immature T1 legs at 0 hours APF expressing *bi>GFP* (grey), immunostained for Dll (red), Dac (green), and Twi (purple). **(F1)** is a maximum projection; **(F2)** is a confocal section; **(F3)** is a maximum projection; **(F4)** shows an enlargement of the dotted box in **(F3)**. Note: bi is a good marker for cluster 14_2 and weakly expressed in cluster 15 **(see** Fig. 4**M****1)**. GFP expression confirms low levels in Twi^−^ cells in the tarsus (Ta), tibia (Ti), and femur, and high expression in Twi^+^ cells in the tibia. **(G1–G4)** Immature T1 legs at 0 hours APF expressing *RtGEF>GFP* (grey), immunostained for Dll (red), Dac (green), and Twi (purple). **(G1)** is a maximum projection; **(G2)** is a confocal section; **(G3)** is a maximum projection; **(G4)** shows an enlargement of the dotted box in **(G3)**. Note: *RtGEF>GFP* is a specific marker for cluster 6, which is localized in the dorsal femur (Fe). **(H1–H3)** Immature T1 legs at 0 hours APF expressing *RtGEF>GFP* (grey), immunostained for **Doc2** (a specific marker of cluster 6, red), and counterstained with DAPI (blue). **(H1)** is a maximum projection; **(H2–H3)** are confocal two different sections showing enlargements of the dotted box in **(H1)**.

**Figure S6.**
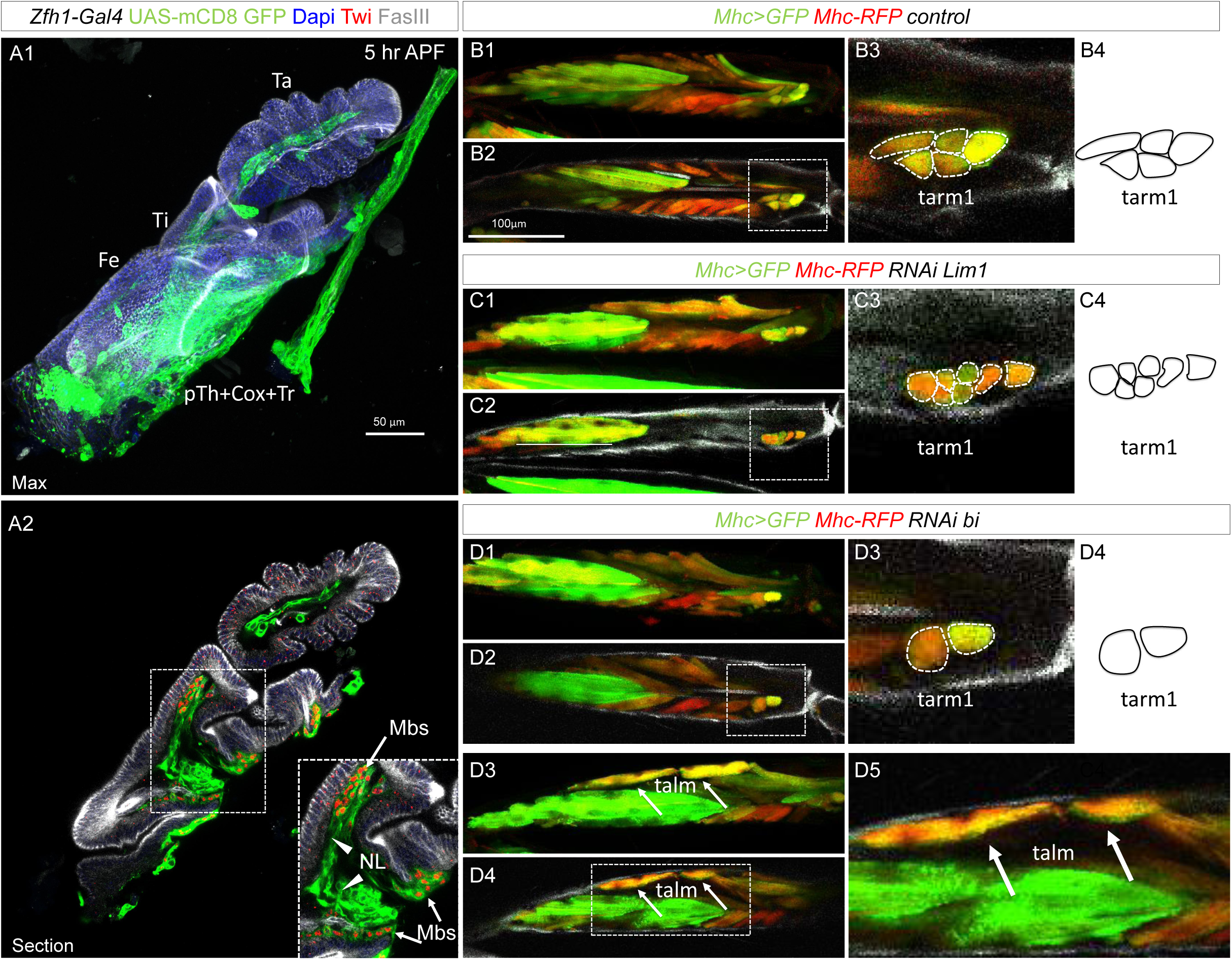
Expression of *Zfh1-Gal4* in immature legs and the effects of *bi* and *lim1* RNAi on muscle architecture using the Zfh1-Gal4 driver. **(A1–A2)** Immature legs at 5 hours APF genetically labeled with *Zfh1-Gal4 > UAS-mCD8::GFP* (green) and immunostained for Twist (Twi, red) and FasIII (gray), with nuclei labeled by DAPI (blue). **(A1)** shows a maximum projection and **(A2)** a single confocal section. The dotted box in **(A2)** highlights Zfh1>GFP+ myoblasts (Mbs, arrows) expressing Twi. Zfh1>GFP+ but Twi– cells correspond to neuronal lamella cells (NL, arrowhead). **(B1–D5)** Adult tibia of T1 legs genetically labeled with *MHC-RFP* (red) and *UAS-mCD8::GFP* (green) under the control of *MHC-Gal4*. Panels show: a control fly **(B1–B4)**, a fly with *bi* knockdown (KD) in MPs, myoblasts, and muscles **(C1–C4)**, and a fly with *Lim1* KD in MPs, myoblasts, and muscles **(D1– D5)**. **(B1, C1, D1, D3)** are maximum projections; **(B2, C2, D2, D4)** are confocal sections; **(B3, C3, D3, D5)** are enlargements of the dotted boxes in **(B2, C2, D2, D4)**; and **(B4, C4, D4)** are schematics of the tarsal muscles. **Note:** In **(B1–D4)**, the number of tarsal muscle fibers is reduced in flies with *bi* KD and increased in flies with *lim1* KD. In **(D3–D5)**, rounded fibers are observed in tarsal muscles of flies with *bi* KD (arrows).

**Figure S7.**
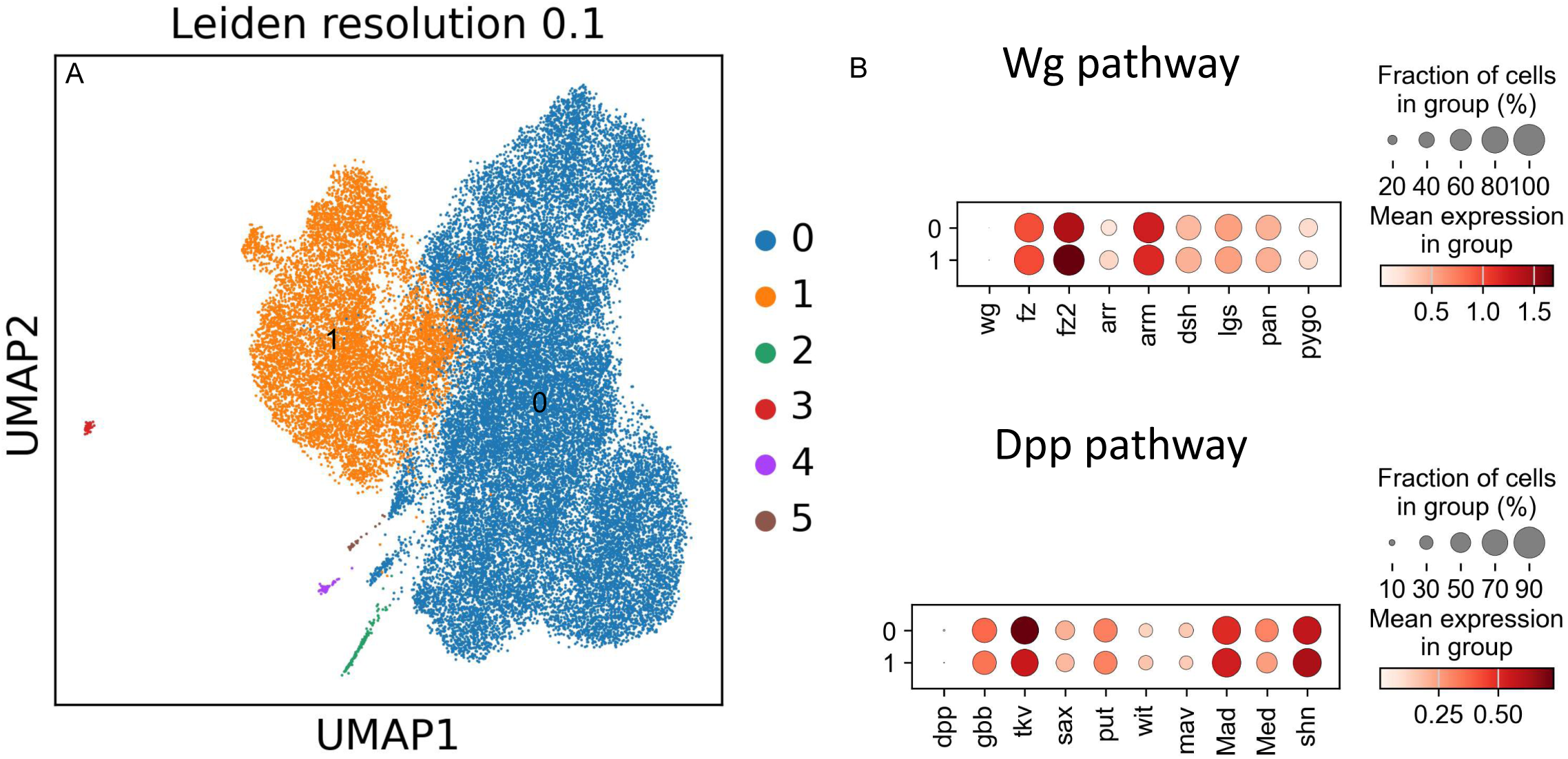
MPs and myoblasts express all components required for Wg/Dpp signal reception and transduction but do not express the morphogens themselves. **(E2, F2, G2)** are maximum projections; **(E3, F3, G3)** are confocal sections; **(E4, F4, G4)** are schematics of the tibia showing the talm, tadm, and their corresponding tendons, talt (cyan) and tadt (pink); the long tendon (lt) is also represented (purple). Cross sections on the right are indicated by dotted lines on the left. **(A)** Low-resolution UMAP revealing two major clusters. **(B)** Dot plot showing the expression of Wg and Dpp, as well as the proteins involved in signal reception and transduction

